# Systematic comparison of Generative AI-Protein Models reveals fundamental differences between structural and sequence-based approaches

**DOI:** 10.1101/2025.03.23.644844

**Authors:** Alexander J Barnett, KC Rajendra, Pratikshya Pandey, Pamodha Somasiri, Kirsten A Fairfax, Sandy Hung, Alex W Hewitt

## Abstract

Recent advances in artificial intelligence have led to the development of generative models for *de novo* protein design. We compared 13 state-of-the-art generative protein models, assessing their ability to produce feasible, diverse, and novel protein monomers. Structural diffusion models generally create designs with higher confidence in predicted structures and more biologically plausible energy distributions, but exhibit limited diversity and strong sequence biases. Conversely, protein language models generate more diverse and novel designs but with lower structural confidence. We also evaluated these models’ ability to generate unique proteins, conditionally based on the Tobacco Etch Virus (TEV) protease. Generative models were successful in producing functional enzymes, albeit with diminished activity compared to the wildtype TEV. Our systematic benchmarking provides a foundation for evaluating and selecting generative protein models, while highlighting the complementary strengths of different generative paradigms. This framework will facilitate an informed application of these tools for bio-medical engineering and design.

## Main

The remarkable versatility displayed by proteins in biological systems makes them an obvious class of interest for *de novo* molecular design. Protein functional properties are largely the product of their ordered tertiary structure, which emerge through folding of the cognate amino acid chain. Advances in deep learning have driven rapid performance gains in the prediction of protein structures and functional properties from arbitrary amino acid sequences^1,2^. Often outperforming traditional biophysical approaches, applications such as Alphafold^3–5^ highlight the capacity of data-driven deep learning models to attain fundamental insights into the relationships between protein sequence, structure, and function. Although predictive models can economically facilitate the assessment of candidate protein sequences, the space of possible sequences is vast, making the ability to appropriately prioritise or sample new proteins essential.

Seeking to leverage the understanding of protein biochemistry uncovered by predictive-AI models, a range of deep generative models have recently been trained on various empirical protein data, including sequences^6–10^, backbone conformations^11–15^, and full tertiary structures^16–18^. During training, these models learn a probability distribution over polypeptide-space, which reflects both the physical constraints imposed by three-dimensional structures, as well as representing the evolutionary pressures present in biological systems. Sampling from this distribution distils polypeptides to a more tractable candidate-space enriched for biologically relevant proteins, and a subset of models can impose specific constraints during training or inference to refine the candidate-space to one enriched for particular functional properties; however, it is currently not clear which generative models perform the best.

Evaluating the capabilities of such generative models is a complex task and little comparative work has been undertaken to systematically assess the merits and shortcomings of different approaches. While it is a reasonable assumption that a model trained on experimentally resolved structures would receive a higher concentration of exploitable information, it is unclear whether this is sufficient to offset the considerably lower volume of data compared to the abundance of known protein sequences. Until recently, most generative models of protein structures have limited themselves to the task of generating candidate backbone conformations, relying on downstream models for sequence design and structural prediction. While more efficient during training, the extent to which the omission of side-chain influences during generation impacts the quality, diversity, and novelty of their designs is not well understood.

Herein, we compare the ability of 13 generative protein models spanning multiple paradigms (**Table 1**) to efficiently generate feasible, diverse, and novel protein monomers. We further apply five of these models to the relatively complex task of designing a monomer around a set of functional motifs from the Tobacco Etch Virus (TEV) Protease to assess their ability to generate proteins subject to specific constraints.

**Table 1:** Overview of Generative Protein Models included in this study.

| Model | Paradigm | Output | Training Data Sources |  |  |  |  | Postprocessing Required for All-Atom Designs |  |  | Ability for TEV protease conditional design |
| --- | --- | --- | --- | --- | --- | --- | --- | --- | --- | --- | --- |
|  |  |  | RCSB PDB | SCOPe | CATH | UniRef | BFD | Backbone Refinement (RosettaMinMover) | Sequence Design (ProteinMPNN) | Structure Prediction (OmegaFold) |  |
| RFdiffusion <sup>11</sup> | D(Str) | BC | ✓ |  |  |  |  |  | ✓ | ✓ | ✓ |
| Genie <sup>12</sup> | D(Str) | BC (Cα) | | ✓ | | | | | ✓ | ✓ | |
| ProteinSGM <sup>13</sup> | D(Str) | BC |  |  | ✓ |  |  | ✓ | ✓ | ✓ |  |
| FoldingDiff <sup>14</sup> | D(Str) | BC |  |  | ✓ |  |  |  | ✓ | ✓ |  |
| FrameDiff <sup>15</sup> | D(Str) | BC | ✓ |  |  |  |  |  | ✓ | ✓ |  |
| Chroma <sup>16</sup> | D(Str) | FTS | ✓ |  |  |  |  |  |  |  | ✓ |
| Protpardelle <sup>17</sup> | D(Str) | FTS |  |  | ✓ |  |  |  | * |  | ✓ |
| ProteinGenerator <sup>18</sup> | D(Seq) | FTS | ✓ |  |  |  |  |  |  | ** | ✓ |
| EvoDiff <sup>6</sup> | D(Seq) | Sequences |  |  |  | ✓ |  |  |  | ✓ | ✓ |
| RITA <sup>7</sup> | APLM | Sequences |  |  |  | ✓ |  |  |  | ✓ |  |
| ProGen2 <sup>8</sup> | APLM | Sequences |  |  |  | ✓ | ✓ |  |  | ✓ |  |
| ProtGPT2 <sup>9</sup> | APLM | Sequences |  |  |  | ✓ |  |  |  | ✓ |  |
| ESM-Design <sup>10</sup> | MTPLM | Sequences |  |  |  | ✓ |  |  |  | ✓ |  |

Abbreviations: D(Str), Diffusion (Structure); D(Seq), Diffusion (Sequence); APLM, Autoregressive Protein Language Model; MTPLM, Masked Token Protein Language Model; BC, Backbone conformations; BC (Cα ), Backbone conformations - Cα coordinates only; FTS, Full Tertiary Structures; TEV, Tobacco Etch Virus.
\* Protpardelle uses ProteinMPNN for Sequence Design at each denoising step of it's structure generation
\*\* ProteinGenerator is adapted from the structural prediction network RoseTTAFold to generate sequence-structure pairs from noised protein sequences

## Results

### Unconditional Monomer Generation

#### Implementation on Google Colaboratory

To assess the capabilities and usability of the selected models at a computational power accessible for open and economical use, we implemented each on the Google Colaboratory (Colab) platform according to their standard installation and execution instructions. RFdiffusion and Chroma provide existing public implementations on this platform, while RITA and ProtGPT2 are both easily accessed through the HuggingFace Transformers library. Most other models were straightforward to implement, requiring at most minor configuration changes and management of python environments to successfully perform free generation. However, ProteinSGM required substantial work to implement on Colab due to its dual use CPU and GPU processing and errors encountered when executing as described in the documentation. The majority of models were capable of generating monomers within a length of 14 to 200 residues, and of performing batched generations at a set length. Exceptions were the autoregressive protein language models (Rita, ProGen2, and ProtGPT2), which can only be supplied with a non-specific maximum generation length; FoldingDiff and ProteinSGM, which have limited output length ranges (≤128 and 40-128 residues respectively); and ESM-Design and Genie, which experienced issues with GPU memory overflow when running batched generations on Colab. We found that ESM-Design with the default number of iterations would take ∼26 hours to generate a single length-200 monomer on Colab. To include it in this study, we significantly decreased the number of iterations to reduce its generation times to be comparable to the other protein language models. Rosetta FastDesign and FastRelax were disabled in the backbone refinement step of ProteinSGM for similar reasons.

#### Generation Time and Designability

We generated approximately 3,000 unconstrained all-atom protein monomer designs of between 14 and 200 residues in length using each selected generative model (**Figure 1A-C, Supplementary Figures 1 & 2, Supplementary Table 1**). Across all models, the observed median walltime required to produce an all-atom design within this length-range generally follows an O(n) or O(n^2^) relationship with respect to length. One notable exception is Genie, for which the median generation walltime does not noticeably increase with monomer length. This is likely due to the fact that Genie generates only the Cα atom positions of the backbone, and therefore the complexity of the task scales more slowly than other Structural Diffusion models, which must generate positions for many and increasingly more atoms. Generative models of full tertiary structures and sequences appear to require shorter median walltimes for inference than those generating backbone conformations. However, the median confidence in the final predicted tertiary structures are consistently higher for generative models of backbone conformations (**Supplementary Figures 3 & 4**). While the speed of the structural diffusion models of full tertiary structures may seem counter-intuitive, given that other methods disregard side chains to increase efficiency, Chroma and Protpardelle were both developed with a clear emphasis on optimisation.

**Figure 1.**
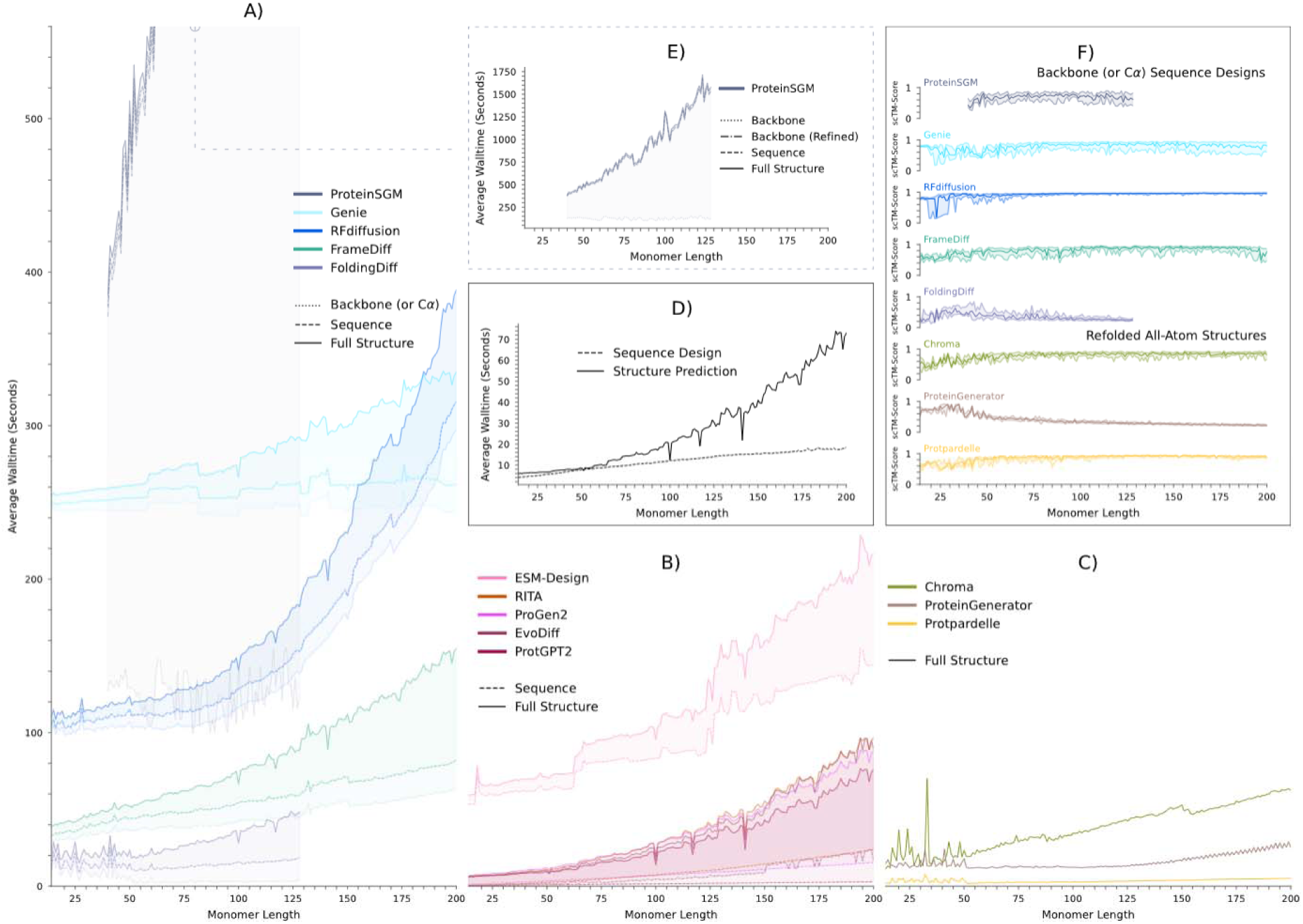
Performance of our selected generative models with respect to median generation walltime and designability. Median walltimes required for unconditional generation of all-atom monomer designs with generative models of A) Backbone Conformations (or Cα positions in the case of Genie), B) Protein Sequences, and C) Full Tertiary Structures, generally increase in a O(n) or O(n^2^) fashion as monomer length increases. D) Post-generation steps of sequence design and structural prediction, where needed, generally represent a small fraction of the total required walltime to produce an all-atom design. E) Expanded view of outlying median walltimes per design for ProteinSGM (largely due to the inclusion of a physical backbone refinement step). F) Agreement between generated backbone conformations (or Cα positions in the case of Genie) and putative full tertiary structures, and between generated full tertiary structures and structures predicted by OmegaFold for their primary sequence (scTM-Score approaching 1 indicates strong agreement). Centre lines represent the median value per-length. Outer lines indicate the two inner quartiles.

The additional wall time required to conduct sequence design (ProteinMPNN^19,20^) and structure prediction (OmegaFold^21^), as required by many models, was not a limiting factor in this length-range compared to initial model inference (**Figure 1D**). The exception is ProteinSGM, which requires an additional, time-intensive step of backbone refinement using Rosetta MinMover^22,23^ (**Figure 1E**). However, this is only the case for a single use of these tools. In reality, most protein design approaches perform multiple iterations of sequence design or structural prediction before selecting the best candidate. Conversely, while a single ProteinSGM generation is significantly slowed by the use of Rosetta MinMover, this can be offset over multiple generations through CPU batch processing.

The designability of generated backbone conformations was assessed by the agreement between the backbone conformation and the final full tertiary structure. We generated 10 putative sequences for each backbone output, and calculated the self consistency TM-Score^24^ (scTM-score) between their predicted structures and the generated backbone (**Figure 1F**). The designability of the backbone conformations appear to be inversely related to the median generation wall-time. RFdiffusion consistently produces final designs with the highest median confidence in the predicted structure and agreement with the generated backbone (**Supplementary Figures 3 & 4**), but also exhibits the poorest scaling of generation time with respect to monomer length.

Designability is consistently weaker for shorter monomers, becoming more stable at protein lengths more commonly observed in nature. Similar results were observed with respect to scRMSD (**Supplementary Figure 3**). We also calculated scTM-scores between the outputs of our three generative models of full tertiary structures and a structure predicted for each of their primary sequences by OmegaFold (**Figure 1F**). Chroma and Protpardelle display similar performance to the more performant backbone models in this metric, while consistently low agreement between generated and refolded structures were observed for ProteinGenerator.

#### Feasibility

We surveyed the sequence content and the geometric and energetic qualities of our generated monomers in comparison to a subset of experimentally resolved protein chains from the PDB^25^ (PISCES) using PyRosetta^26^ (**Figure 2, Supplementary Figures 5-10**). Both the experimentally resolved chains and our generated designs exhibit largely uniform amino acid preferences with respect to residue position (**Supplementary Figure 7**), but diverging patterns were observed in the overall enrichment for different amino acids throughout their sequences. EvoDiff and the three autoregressive language models appear to best capture the natural distribution of amino acid preferences while ESM-Design monomers exhibit an unusual uniform preference for all residues except cysteine. A consistent enrichment in lysine and glutamic acid, and depletion of asparagine, glutamine, histidine, tryptophan and serine is observed in monomers generated by structural diffusion models.

**Figure 2.**
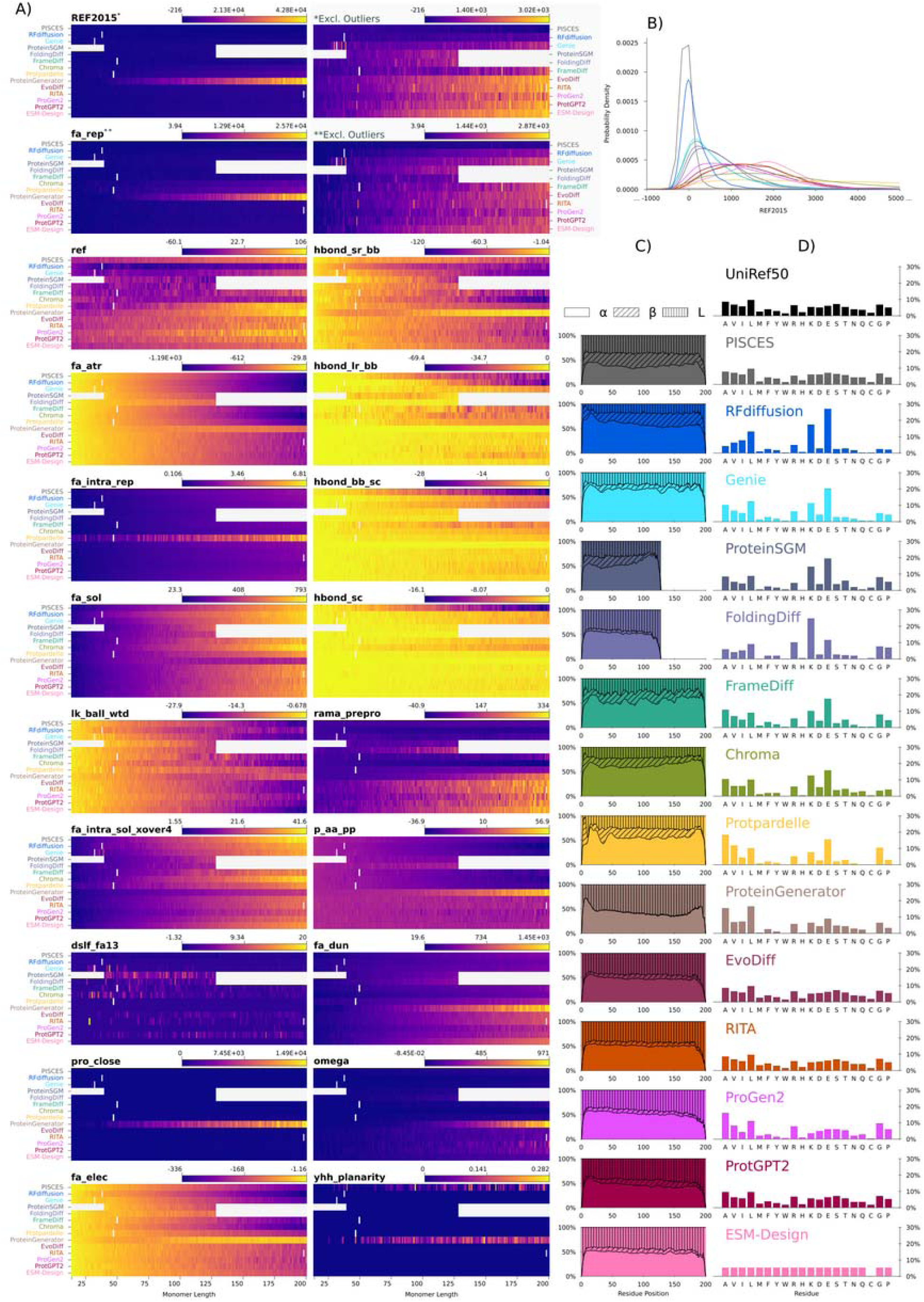
Biochemical profile of Generative AI-proteins. A) Rosetta Energy Score (REF2015) and subterm values for our generated monomers and a subset of resolved protein chains from the PDB (PISCES) vs monomer length. B) Probability density estimation (KDE) of REF2015 values from our generated monomers and PISCES chains. C) Percent (%) enrichment for alpha helix, beta strand and coil secondary structures per residue position for our generated monomers and PISCES chains. D) Overall enrichment of each amino acid in our generated monomers, PISCES chains, and the UniRef50 dataset.

I--strands were often under-represented in the secondary structures of generated designs compared to the experimentally resolved chains, particularly those from generative models of sequences. RFdiffusion, FrameDiff, Chroma and Protpardelle appear to best capture the natural distribution of secondary structure motifs with respect to residue position. With the exception of ProteinGenerator, all models were able to recreate the distributions of Φ, Ψ, and ω angles observed in experimentally resolve chains (**Supplementary Figure 8**). All models generate structures with narrower distributions of N-Cα, Cα-C, and C-O bond lengths than observed in the experimentally resolved chains with the exception of Protpardelle, where the distributions were far broader than those observed for naturally occurring proteins. With the exception of Chroma, all models generate structures with a broader range of C-N bond lengths than observed in the experimentally resolved chains (**Supplementary Figure 9**). Designs from ProteinGenerator often exhibit irregular χ angle distributions in their side chains (**Supplementary Figure 10**).

With the exception of RFdiffusion, all models generate monomers with much higher total rosetta energies^27,28^ (REF2015) than those observed in the experimentally resolved chains. This is largely a consequence of high Lennard-Jones repulsion values between atoms in different residues (fa_rep). Designs from ProteinGenerator and the other generative models of full tertiary structures exhibit particularly high values for fa_rep; however, this is less prominent in the structures predicted for their primary sequences by Omegafold (**Supplementary Figure 6**).

The distributions in the energetic sub-terms of REF2015 vary between different generative models, as well as between generated designs and experimentally resolved chains. Consistent with our observations in overall amino-acid enrichment, designs generated by structural diffusion models differ from the experimentally resolved chains in reference energies for each amino acid (ref). Protpardelle designs exhibit higher Lenard-Jones repulsion between atoms in the same residue (fa_intra_rep) than other models and experimentally resolved chains. With the exception of FoldingDiff and, in some cases, Protpardelle, structural diffusion models consistently differ from sequence-based models in Lennard-Jones attraction between atoms in different residues (fa_atr), inter-residue Lazaridis-Karplus solvation energies (fa_sol), coulombic electrostatic potentials (fa_elec), short-range backbone hydrogen bond energies (hbond_sr_bb), backbone torsion preferences (rama_prepro), penalties for deviant dihedral angles (omega), and backbone-dependent probabilities of amino acid identity (pp_aa_pp). Structural diffusion appears to give more realistic results for fa_atr, fa_elec, rama_prepro, omega, and p_a_pp; while sequence based models give more biologically plausible values for fa_sol and hbond_sr_bb. Nearly all models struggle to recreate reference distributions for sidechain-backbone and sidechain-sidechain hydrogen bonding energies (hbond_bb_sc, hbond_sc), sidechain hydroxyl group torsion preferences (yhh_planarity), disulfide bridge energies (dslf_fa13), and long-range backbone hydrogen bond energies (hbond_lr_bb).

#### Diversity and Novelty

To explore the distribution of generated monomers throughout the protein universe, we embedded their sequences, the sequences of the PISCES chains, and a 1% subsample of the UniRef50 dataset^29^ in the ESM large language model^30^, and performed t-SNE dimensionality reduction^31,32^ on the embedding vector to two dimensions (**Figure 3**). Chains from the PISCES set populate, in a length dependent manner, a similar overall region of this 2D space as the UniRef50 sequences, although some specific high density clusters are less apparent (**Supplementary Figure 11**). We also assessed the structural diversity of our generated designs by quantifying the number of structural clusters detected by MaxCluster^33^, and their novelty, by querying each against the ESMAtlas30 database^30^ using Foldseek^34^. The uniform distribution of cluster labels and TM-scores observed in our projections (**Supplementary Figures 12 & 13**) suggests that distances in this space are likely more indicative of sequence differences than structural ones.

**Figure 3.**
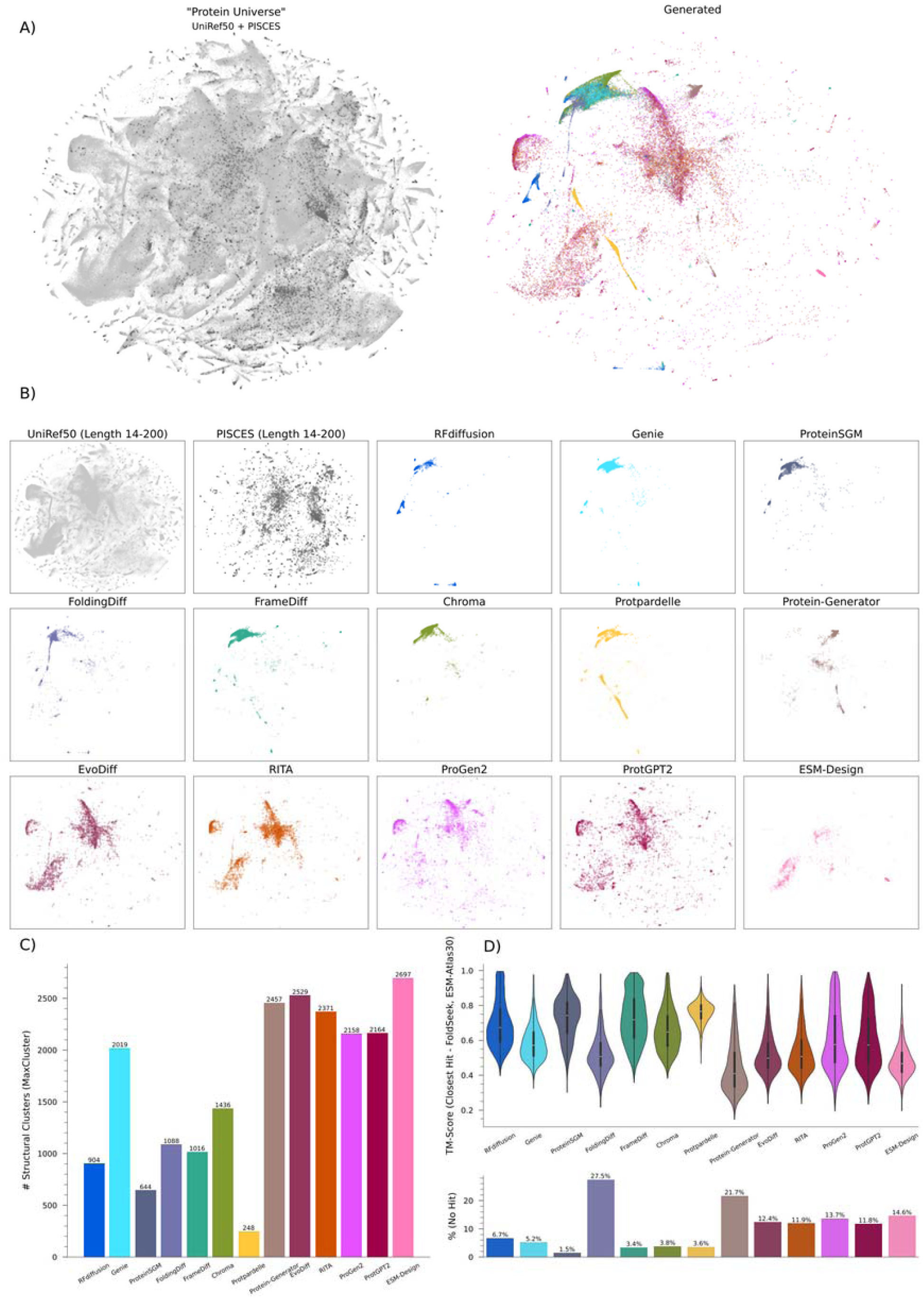
Diversity of naturally occurring and generative AI-proteins. A) *t-SNE* visualisations of the “Protein Universe” — the sequences of a 1% subsample of UniRef50 and our experimentally resolved protein chains (PISCES), along with our generated monomers — embedded in the ESM Large Language Model. B) *t-SNE* visualisations of sequences between 14-200 residues, separated by source. C) Counts of structural clusters detected in the pool of length 14-200 generated monomers for each of our models. D) Distribution of TM-Scores for the best hits of our generated monomers when queried against the ESMAtlas30 database using Foldseek (top), % of generated designs per model for which no hits were returned (bottom).

Similar patterns in distribution throughout this space can be observed between generative models of the same paradigm. Monomers generated through structural diffusion appear to only occupy a small region in comparison to both the UniRef50 and PISCES sequences, whereas generative models of sequences appear to more evenly populate the space of similar length natural proteins. Dispersion of ProteinGenerator, and ESM-Design monomers lie somewhere between the two. Monomers generated by RFdiffusion are the most highly localised within this space, while those from ProtGPT2 have a distribution most similar to that seen in the UniRef50 dataset. Designs from sequence-based models also displayed a larger total number of structural clusters than structural diffusion methods, with the exception of Genie, whilst designs from Protpardlle exhibit the least structural variety. RFdiffusion and FrameDiff produce a more even distribution of cluster sizes compared to the other structural diffusion models (**Supplementary Figure 14**). Generative models of sequences also have a higher tendency toward structural novelty in their designs, with lower TM-Scores for their closest Foldseek query hits and a higher frequency of queries with zero hits. The exception is FoldingDiff, which displays a similar level of structural novelty in its designs to generative models of sequences and the highest incidence of queries with zero hits. Similar results are observed when considering all hits (**Supplementary Figure 15**).

### Conditional design around a set of functional motifs

#### Conditional generation of TEV Protease designs

RFdiffusion, Chroma, Protpardelle, ProteinGenerator, and EvoDiff were selected to redesign the Tobacco Etch Virus (TEV) protease by performing conditional generation of a 237 residue monomer around five fixed motif regions. Genie, FoldingDiff, FrameDiff and ESM-Design do not currently support conditional generation, while ProteinSGM cannot generate monomers of sufficient length. Autoregressive protein language models are only capable of conditional design around a fixed motif at the beginning of a sequence.

Conditional designs by ProteinGenerator were produced using two approaches: specifying only the motif sequences, and specifying the full structural motifs. Similarly, Chroma and RFdiffusion designs were produced by specifying the structural motifs from both the TEV protease monomer in isolation, and in complex with its recognition sequence. Additionally, sequence redesign of the backbones generated by RFdiffusion was conducted with both full flexibility and with motif sequences fixed. As a comparison, we also generated TEV-based protease designs using only ProteinMPNN sequence design with fixed motif sequences. One hundred TEV protease designs were generated for each approach (**Supplementary Table 2**), and the ten sequences for each model with the lowest RMSD to template structures in the motif regions were selected for synthesis and *in vitro* validation.

#### Functional Validation

To compare the catalytic activity, designed monomers were expressed in BHK21 cells together with a tetracycline inducible green fluorescent protein (GFP) and a synthetic protein consisting of tetracycline-controlled transactivator (tTA) tethered via a linker containing the TEV endogenous catalytic site (ENLYFQ’S) to a transmembrane domain protein. The transmembrane domain protein fused tTA is localised to the plasma membrane, and thus the GFP signal is low in the absence of an active TEV protease, but an active protease cleaves tTA enabling its translocation to the nucleus and induction of GFP expression (**Figure 4A**).

**Figure 4.**
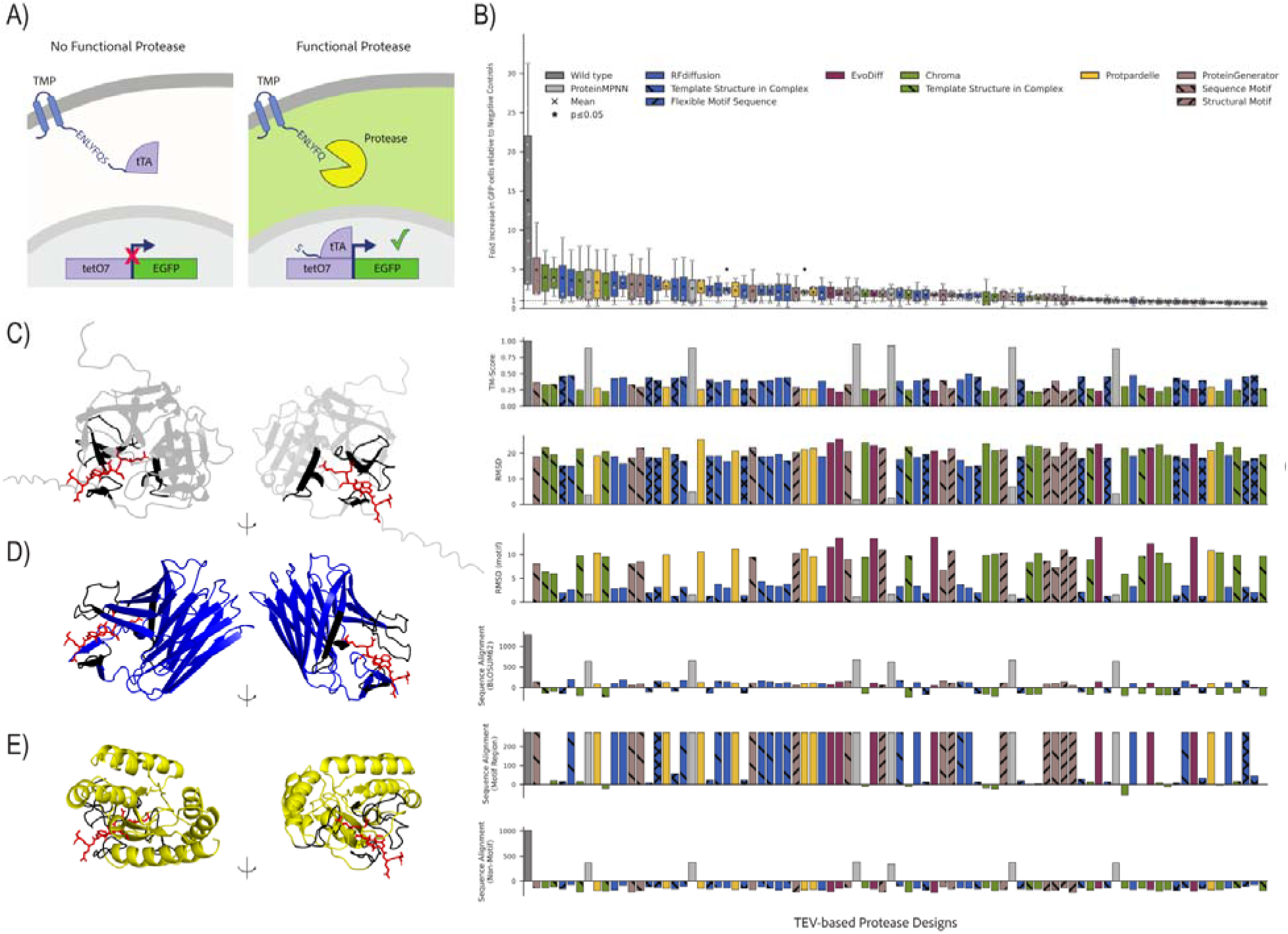
In-vitro validation of conditional designs around a set of functional motifs from the Tobbaco Etch Virus (TEV) Protease. A) *Schematic diagram illustrating tTA localisation inside the cell in the absence and presence of a functional TEV protease. Following cleavage by protease, the tTA translocates to the nucleus and induces the EGFP gene expression.* B) Eighty seven TEV designs generated using 13 different models were ranked based on their enzyme activity (across 3 technical replicates), measured as a fold increase in GFP fluorescence compared to the no TEV control. No trend is apparent between enzyme activity and pairwise structural and sequence alignment scores (TM-Score, RMSD, BLOSUM62) of the designs against the template structure (Lower). C) Template Structure of wild type TEV protease S219V (Grey) used for conditional designs in complex with the recognition sequence (red). D) TEV104, a TEV protease design produced with RFdiffusion and E) TEV26 - a TEV protease design produced with Protpardelle. Functional motifs are highlighted in Black.

Twenty-three of the 110 selected design monomers could not be successfully cloned, potentially as a result of the instability or toxicity of the synthesised sequence towards the host *E.coli* cells. We evaluated the functional activity of 87 TEV designs and ranked them based on their enzyme activity, measured as a fold increase in GFP fluorescence compared to the no TEV control (**Figure 4B**). 34 designs exhibited greater than 2-fold increase in GFP fluorescence, suggesting potential protease activity; however, only two designs (designated TEV104 and TEV26 generated using RFdiffusion and Protpardelle, respectively) were found to have a nominally significant activity (P=0.045 and P=0.023, respectively) compared to the negative controls, though this was not significant following Bonferroni correction. The contact residues within the motif of the wildtype TEV protease (**Figure 4C**) are conserved and in close contact with the target substrate sequence in both TEV104 and TEV26 designs, suggesting critical interactions are preserved (**Figure 4D** & **4E**). Additionally, these residues demonstrate a similar spatial orientation around the target substrate, indicating recapitulation of key interaction motifs necessary for substrate binding and cleavage activity. Interestingly, there was no discernible relationship between the design strategy and protease activity in the tested TEV designs, and the efficiency is still far less compared to the wildtype protease, suggesting the need for further optimisation.

## Discussion

We adopted a rational, quantitative approach to systematically benchmark generative protein models. In initial reports, the qualitative performance of these models has often been assumed from the degree of model fit to training data, or evaluated from a few selected outputs. The lack of a systematic approach has resulted in an incomplete understanding of the merits and shortcomings of current popular approaches, the rationale for their use, or potential complementary strategies. The trends we have observed across our selected metrics for monomer designability, feasibility, diversity, and novelty, have highlighted performance trade-offs between models and revealed fundamental differences between the two popular paradigms of structural diffusion and protein language modelling. Additionally, we have assessed certain practical considerations of running these models, with the goal of functional and accessible *de novo* design.

Models based on structural diffusion generally produce designs with higher confidence in the putative structure, and more biochemically appropriate distributions of energy terms and secondary structure features. However, they are less structurally diverse, and exhibit a strong bias toward particular sequence content that is inconsistent with both data comparable to their training examples (PISCES chains) and the larger space of known protein sequences (UniRef50). While it is possible that this bias towards particular sequence content and limited structures is introduced during sequence design with ProteinMPNN, these results are also observed for Chroma, which uses its own novel sequence design network.

Structural diffusion models appear to have learned to generate a narrow range of outputs, rather than effectively capturing or generalising beyond the distribution of their training data. The inclusion of side-chain influences in the generation process does not appear to improve this limited diversity. Interestingly, the tertiary structures produced by such methods were also less biochemically feasible than the structures predicted for their primary sequences by Omegafold, largely due to poor side-chain conformations. However, considering the short generation times achieved by these methods, incorporating a post-generation ‘refolding’ step is a feasible strategy to overcome this.

Protein language models generally produce a diverse range of designs that are more likely to be novel, but typically have lower confidences in their putative structures, and are potentially less biochemically plausible. Nevertheless, it is less clear whether this represents a higher level of intrinsic disorder for these sequences, or reflects the limitations of the structural prediction method over the more diverse space of sequences explored by these models. The sequence diffusion approach of EvoDiff appears to generally result in similar designs to the autoregressive language models. ProteinGenerator, a model that combines sequence diffusion with a structure-aware objective function, largely produces biochemically unrealistic tertiary structures. ESM-Design also exhibited relatively poor performance, with a near-uniform residue preference observed in its generated sequences. Nevertheless, given the significant reduction in iterations necessary for inclusion, this observation likely reflects the computational power required to use this tool effectively, rather than its inherent capabilities. Similarly, excluding Rosetta FastDesign and FastRelax from the backbone refinement step of ProteinSGM is anticipated to have impacted its optimal performance.

For the five models capable of executing our TEV protease redesign task, we found no significant evidence of increased activity over negative controls or meaningful differentiation between different models or design strategies upon functional validation. This suggests that further improvements are needed before these models, all based on sampling from learned diffusion processes, can complete such tasks without substantial oversight and optimisation. More iterative protein design approaches, such as SCUBA^35^, which continuously samples and optimizes backbone structures using a learned energy function, may be better suited to such tasks.

Even when data distributions are explicitly modelled and likelihoods are tractably computed, these factors alone do not ensure high-quality, diverse proteins or generalisation beyond current training datasets. Many current models generate incomplete representations of proteins, such as sequences or backbone conformations, making the quality and diversity of their designs dependent on the predictive models used alongside them. By directly examining the attributes of their outputs, we have systematically compared these models at the point most relevant to *de novo* design.

## Methods

### Generation of Protein Designs

All protein generations, backbone sequence designs, and structure predictions were performed using a T4 GPU on the Google Colaboratory platform. Sequence designs were conducted on Cα-only backbone structures using ProteinMPNN^19,20^. Ten prospective sequences were generated per backbone at a sampling temperature of 0.1. Structure predictions were conducted with OmegaFold^21^ using default parameters.

### Unconditioned monomer designs

For each of the 13 generative models, a distribution of lengths between 14 and 200 residues (**Supplementary Figure 2)** was sampled using an empirical cumulative distribution function estimated from the repartition of Swiss-Prot sequences by size reported by the Uniprot^29^ statistics portal (Accessed January 2024). Unconditioned monomer designs were generated at these lengths using the default parameters for each model as described in their documentation unless otherwise noted. Designs at each length were batched where possible, with generation wall-time assumed to be evenly distributed between designs. For autoregressive large language models, where only a maximum length can be specified at inference, random length monomers were generated with an appropriate maximum-length parameter until the desired distribution could be subsampled.

Existing Google Colaboratory implementations were available for RFdiffusion and Chroma and were used without alteration beyond the recording of metadata. Minimal adjustments were made in the implementation of other models to enable execution on Google Colab, facilitate batching, manage GPU memory, and reduce model reloading between generations. The Rosetta refinement step for ProteinSGM was conducted on a Google Colaboratory CPU instance, as suggested by its documentation^13^. The Rosetta FastDesign and FastRelax were excluded from this step due to long compute times. Generations with ESM-Design and Genie were unable to be batched due to high GPU memory usage. ESM-Design was run with a significantly reduced number of iterations, 250 reduced from the default 170000, to lower its generation wall-time to be comparable to other models.

### Conditional Design around defined Tobacco Etch Virus (TEV) Protease motifs

#### Definition of Function Motifs and Creation of Template Structures

A structure was predicted for the TEV protease S219V sequence reported by Packer et al.^36^ in complex with its recognition sequence using AlphaFold^3^ (ColabFold^37^) and used to create template structures. Interface residues between the two chains were identified in PyMol^38^ using a script by Jason Vertrees^39^, and used to define five fixed motif regions to supply to the generative models (RES 28-33, 47-51, 140-152, 168-179, 212-221).

#### Generation of conditioned monomers and complexes

Motif regions were provided to each model for conditional generation as outlined in their documentation. Sequence design was conducted on the RFdiffusion backbones with ProteinMPNN defaults and with an overwhelming bias provided towards the motif sequence residues in the motif region. Ten putative sequences were designed for each backbone, and the sequence with the structure most consistent with the backbone (highest scTM-score^24^) was selected. The biassed ProteinMPNN approach was also applied directly to the template structure as a comparison.

#### Selection of Designs

Ten TEV scaffold designs were selected for each conditional design approach. Designs were filtered to those in the bottom quartile of Rosetta Energy^27,28^ (REF2015) scores and the ten designs with the lowest RMSD to template structures in the motif regions were selected (**Supplementary Table 2**). The tertiary structures of Chroma, Protpardelle, and ProteinGenerator designs were refolded with OmegaFold^21^ before selection, due to high observed REF2015 scores for their initial outputs.

### In-silico Analysis

All computational analyses of designs were performed on the Tasmanian Partnership for Advanced Computing (TPAC) Rosalind HPC cluster using multiple Intel(R) Xeon(R) CPU E5-2680 v4 CPUs.

The PISCES server^25^ was used to create a comparison dataset of protein chains from the PDB fulfilling the following criteria: determined by X-ray-diffraction, maximum pairwise sequence identity = 25, Minimum resolution = 0.0, Maximum resolution = 2.0, Maximum R-value (X-ray) = 0.25, Minimum chain length = 40, maximum chain length = 10000, including chains with breaks and missing residues due to disorder. Chain .pdb files were cleaned using pyrosetta.toolbox.cleaning.cleanATOM()^26^.

Outputs from Chroma in .CIF format were converted to .PDB format using PyMOL^38^.

#### Designability

Folded structures for backbone sequence designs and ‘refolds’ of full tertiary structure outputs were aligned to their respective backbone/original design and scTM-score^24^ and scRMSD calculated using the *tmtools* python package and Biopython’s *SVDSuperimposer* module respectively. The backbone sequence design with the highest scTM-score was chosen to represent the design in all further analyses (**Supplementary Table 1**). scRMSD calculations between all-atom designs and refolds were performed on an all-atom basis.

#### 2D Visualisation

The mean embeddings over the sequences of all unconditioned monomer designs, PISCES chains and a 1% subsample of the UniRef50 dataset were extracted from the final layer of the ESM-2 language model^30^ (*esm2_t33_650M_UR50D*) using the *esm-extract* cmd line interface. These 280-dimensional representations were further reduced to 2D coordinates for visualisation using t-SNE-CUDA^31,32^.

#### Novelty

Unconditioned monomer designs were queried against the ESMAtlas30 dataset^30^ to retrieve TM-score normalised by query/target length, average alignment LDDT and probability of homology using Foldseek^34^.

#### Diversity

For each generative model, structural clusters were assigned to unconditioned monomers at each length with MaxCluster^33^ using sequence-independent, average-linkage clustering: *maxcluster64bit -l <pdb_list> -C 2 -in -R1 ./tm_results.txt -Tm 0.5*.

#### Feasibility

Phi (Φ), Psi (Ψ), Omega (Ω), and Chi (χ) angles; backbone bond lengths; sequence; secondary structure annotation; and Rosetta REF2015 energy terms^27,28^ were calculated for each final monomer design using PyRosetta^26^.

### In vitro Analysis

#### Functional Assessment of Monomeric TEV Designs

##### DNA Cloning

A stable and efficient TEV mutant (S219V)^36,40^ and synthetic DNA sequences encoding the TEV designs were codon-optimised for expression in Baby Hamster Kidney (BHK) cells and were purchased as eBlock gene fragments from Integrated DNA Technologies. The gene fragments were cloned into a mammalian expression vector (pCMV-phenomycin) using NEBuilder HIFI DNA Assembly Master Mix (E2621L; New England Biolabs). The assembled plasmids were transformed to NEB Stable Competent *E.coli* grown in Luria broth or agar media supplemented with 100ug/mL carbenicillin. Plasmids were purified using the Monarch Plasmid Miniprep Kit (New England Biolabs).

##### *In vitro* TEV *activity assay*

A tetracycline-controlled transactivator (tTA)-based fluorescent reporter system was used to determine the protease activity of the TEV designs^41^. The reporter system comprised of two components, a tTA fused to a transmembrane domain via a peptide linker containing an endogenous TEV cleavage site (ENLYFQ’S) and an EGFP, expressed under the control of a CMV (CMV-RHO2-TEVcs-tTA) and a seven repeat Tet operator (TETO7-EGFP), respectively.

The reporter and TEV design plasmids were co-transfected into BHK cells using TransIT-X2 following the manufacturer’s protocol. Cells were seeded onto a 48-well cell culture plate at a density of 2.5x10^4^ cells per well, a day prior to transfection. A transfection complex was prepared with 100 ng of each CMV-TEV design, CMV-RHO-TEVcs-tTA, and TETO7-EGFP plasmids, 30 uL optiMEM, and 0.9 uL TransIT-X2. CMV-TEV designs were randomly assigned in triplicate across twelve plates, with each plate containing a positive (wildtype TEV) and negative control (no TEV). The complex was incubated for 20 min at room temperature before adding to the cells. Edge cells contained media only, to avoid edge effects. Twenty-four hours later, EGFP expression was measured using a Carl Zeiss™ Axio Vert.A1 Inverted Microscope. Images were acquired with a 5X objective lens. The same field-of-view was used for the corresponding brightfield image, with auto-exposure settings used for image capture.

##### *Image* Analysis

ilastik^42^, a machine-learning tool for bioimage analysis, was used to quantify EGFP expression using the Cell Density Counting module. The following ‘Features’ were selected from the Cell Density Counting interface: for GFP analysis Color/Intensity (Gaussian smoothing): 0.30, 0,70. 1.00, 1.60, 3.50, 5.00 and 10.00 for σ0, σ1, σ2, σ3, σ4, σ5, and σ6, respectively; for BF analysis Edge (Laplacian of Gaussian): 0.70, 1.00, 1.60, 3.50, and 5.00 for σ1, σ2, σ3, σ4 and σ5, respectively. Under the ‘Counting’ interface, a RandomForest algorithm was used (Ntrees: 10, and MaxDepth: 50), with a sigma value of 8 and 2.5 for BF and GFP analyses, respectively.

Separate models were trained for each replicate for EGFP images and brightfield images. Raw images (.jpeg format) were uploaded to ilastik, and EGFP expressing cells were demarcated in the ‘Foreground’, whereas the ‘Background’ was indicated by areas devoid of a EGFP signal. For cell density counting for bright field images, individual foci were indicated as ‘Foreground’ in areas in which cell surface area was greatest, whereas ‘Background’ was indicated using a fine ‘1’ setting ‘size’ for the intervening spaces of cells in images that had adequately sparse density.

To account for the differences in total cell population, the number of EGFP expressing cells was normalised to the number of cells observed in the brightfield. EGFP expression in the TEV design group was compared to the no TEV control group, and the fold increase was calculated by dividing the number of normalised EGFP positive cells in the TEV design group by that of the negative (no TEV) control group. Statistical analysis was performed using a t-test.

## Code availability

Code for implementing each model on google colab, and analysing the generated data are available at: https://github.com/hewittlab/Systematic-comparison-of-Generative-AI-Protein-Models

## Data availability

The main data supporting the results in this study are available within the paper and its Supplementary Information.

## Acknowledgements

This work was supported by an Australian National Health and Medical Research Council Leadership Award (A.W.H.). We would like to acknowledge the use of the high performance computing facilities provided by Digital Research Services, IT Services at the University of Tasmania.

## Author contributions

A.J.B, R.K. and A.W.H. designed this study. A.J.B. implemented all models and performed the computational benchmarking. R.K performed the in vitro analysis with assistance from P.P., P.S., K.A.F., and S.H. A.W.H. supervised the work. All authors contributed to the writing of the paper.

