## Supplementary for "Systematic comparison of Generative AI-Protein Models reveals fundamental differences between structural and sequence-based approaches"

**Supplementary Information**

**Systematic comparison of Generative AI-Protein Models.**

**Supplementary Figures:**

**Supplementary Figure 1. Number of unconstrained all-atom protein monomer designs of between 14 and 200 residues in length generated per each generative model.**

**Supplementary Figure 2. Distribution of monomer lengths generated with selected models.**

**Supplementary Figure 3. Agreement between putative tertiary structures/refolds and generated backbones/generated tertiary structures (scTM-Score, scRMSD), and median OmegaFold structure prediction confidence vs generated monomer length.**

**Supplementary Figure 4. Median Omegafold structure prediction confidence for final selected backbone sequence designs and generated sequences vs generated monomer length.**

**Supplementary Figure 6. Rosetta Energy Scores (REF2015), subterm values, and secondary structure enrichment vs monomer length/position for PISCES chains and the structures predicted by OmegaFold for the primary sequences of monomer designs from Chroma, Protpardelle, and ProteinGenerator.**

**Supplementary Figure 7. % Amino acid enrichment per position for our generated monomers and PISCES chains.**

**Supplementary Figure 8. Distributions (KDE probability density estimation) of phi (Φ), psi (Ψ), and omega (ω) dihedral angles observed in our generated monomers and PISCES chains.**

**Supplementary Figure 9. Distributions (KDE probability density estimation) of N-Cα, Cα-C, and C-O bond lengths observed in our generated monomers and PISCES chains.**

**Supplementary Figure 10. Distributions (KDE probability density estimation) of side-chain torsion angles ( χ1, χ2, χ3, χ4) observed in our generated monomers and PISCES chains for each residue type (i-XX).**

**Supplementary Figure 11. Distributions of sequence lengths of a 1% subsample of UniRef50, PISCES chains, and our generated monomers, throughout the t-SNE visualisations of their embedding-vectors in the ESM Large Language Model.**

**Supplementary Figure 12. Distributions of the best-hit TM-Scores for our generated monomers when queried against the ESMAtlas30 database in the t-SNE visualisations of their embedding-vectors in the ESM Large Language Model.**

**Supplementary Figure 13. Distributions of the structural cluster labels identified by MaxCluster for our pools of generated monomers in the t-SNE visualisations of their embedding-vectors in the ESM Large Language Model.**

**Supplementary Figure 14. Distributions of sizes for structural clusters identified by MaxCluster for our pools of generated monomers.**

**Supplementary Figure 15. Distributions of TM-Scores of our generated monomers, by model, when queried against the ESMAtlas30 database.**

**Supplementary Tables:**

**Supplementary Table 1. Protein monomer designs generated between 14 and 200 residues in length using each selected generative model.**

**Supplementary Table 2. Redesign of the Tobacco Etch Virus (TEV) protease by performing conditional generation of a 237 residue monomer around fixed motif regions.**

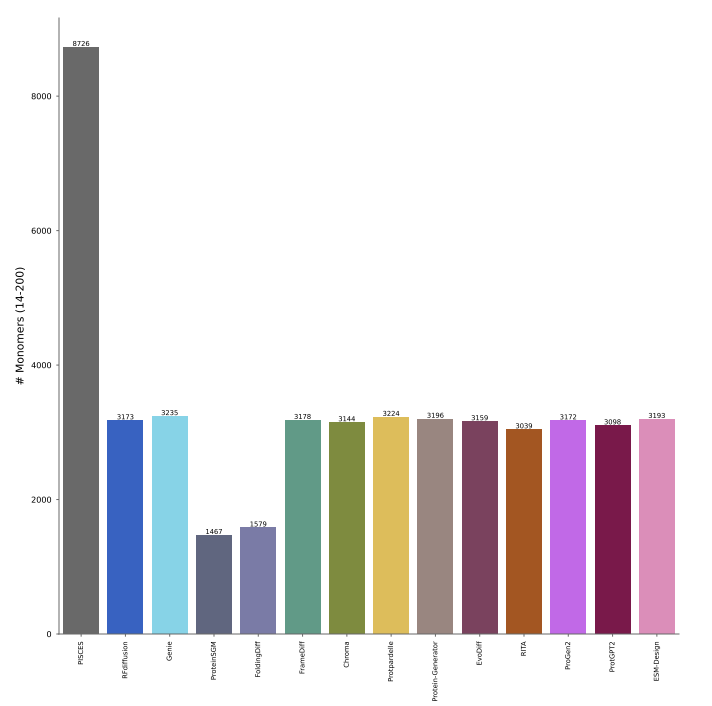

### **Supplementary Figure 1. *Number of unconstrained all-atom protein monomer designs of between 14 and 200 residues in length generated per each generative model.***

*FoldingDiff and ProteinSGM were unable to generate monomers outside of their limited output length ranges (≤128 and 40-128 residues respectively). Number of chains in the PISCES set of experimentally resolved protein chains provided for comparison.*

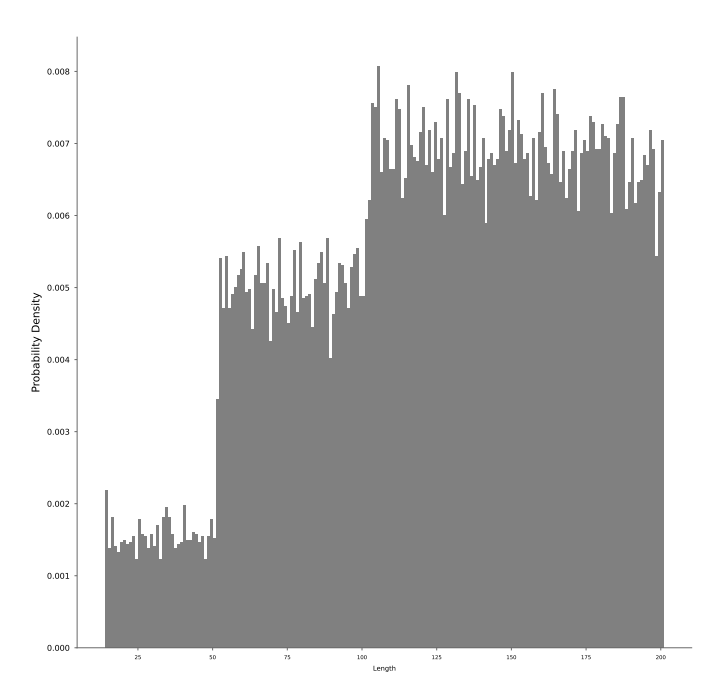

### **Supplementary Figure 2. *Distribution of monomer lengths generated with selected models.***

*Distribution was modelled on the repartition of Swiss-Prot sequences by size as reported by the Uniprot statistics portal (Accessed January 2024).*

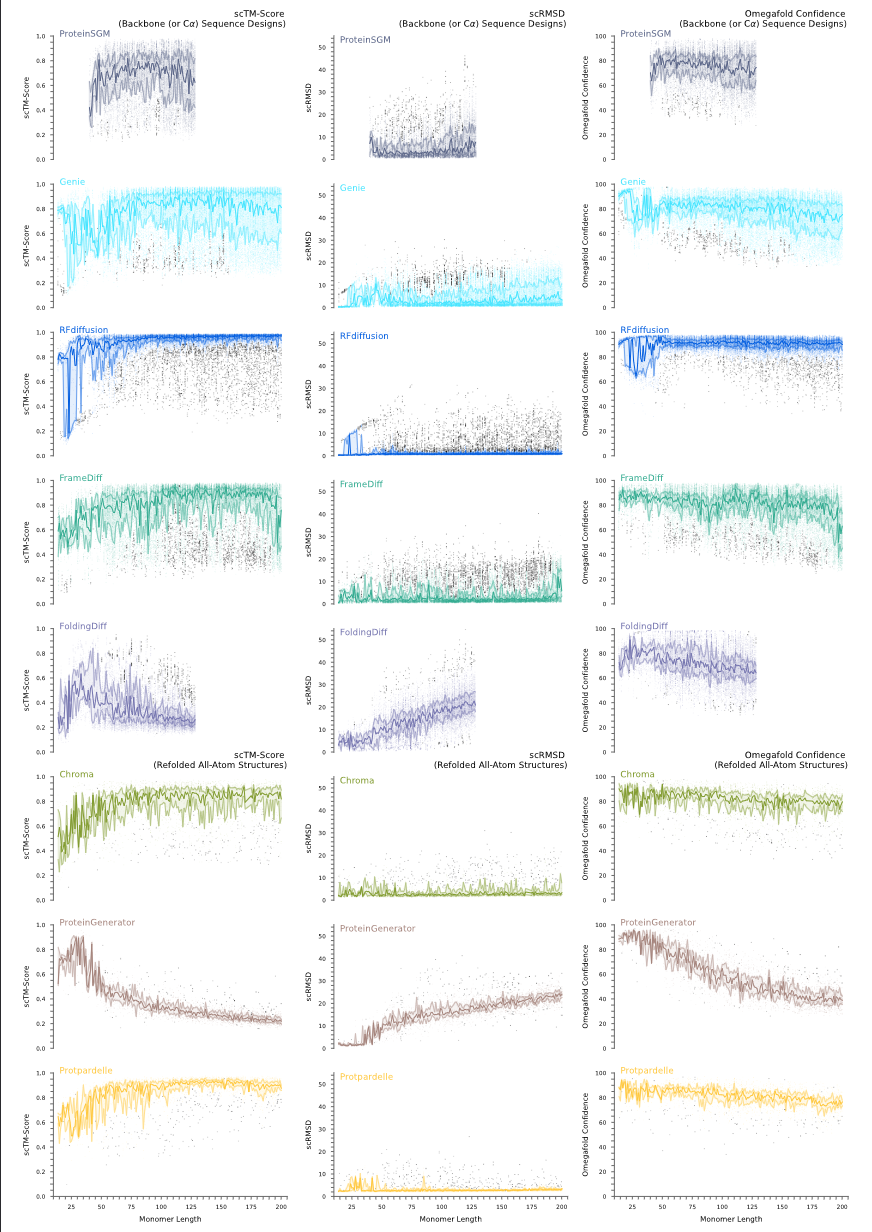

### **Supplementary Figure 3. *Agreement between putative tertiary structures/refolds and generated backbones/generated tertiary structures (scTM-Score, scRMSD), and median OmegaFold structure prediction confidence vs generated monomer length.***

*Centre lines represent the median value per-length. Outer lines define the two inner quartiles. Raw data points are displayed in the model colour with outliers are displayed in black. In the case of Genie, for which only Cα positions are generated, agreement is between the generated backbone Cα atoms and those of the putative tertiary structure.*

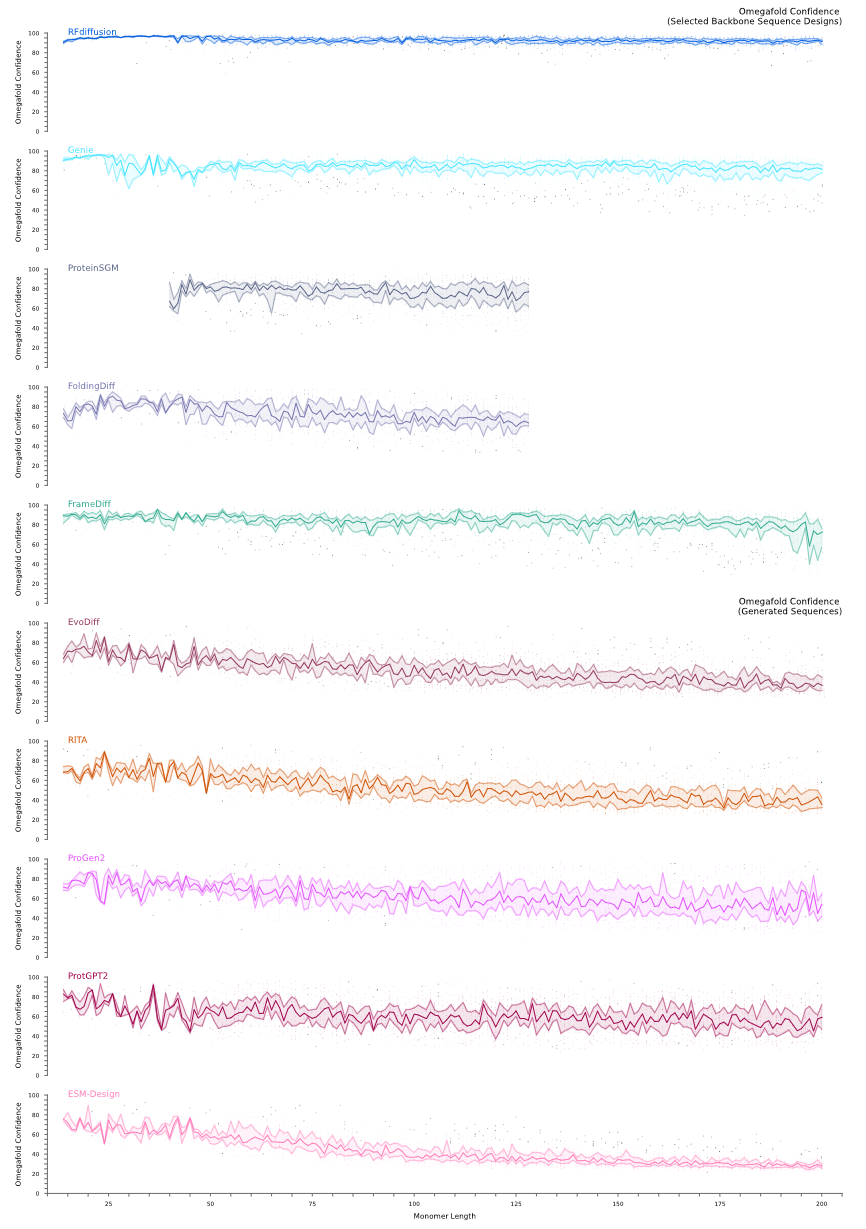

### **Supplementary Figure 4. *Median Omegafold structure prediction confidence for final selected backbone sequence designs and generated sequences vs generated monomer length.***

*Centre lines represent the median value per-length. Outer lines define the two inner quartiles. Raw data points are displayed in the model colour with outliers displayed in black.*

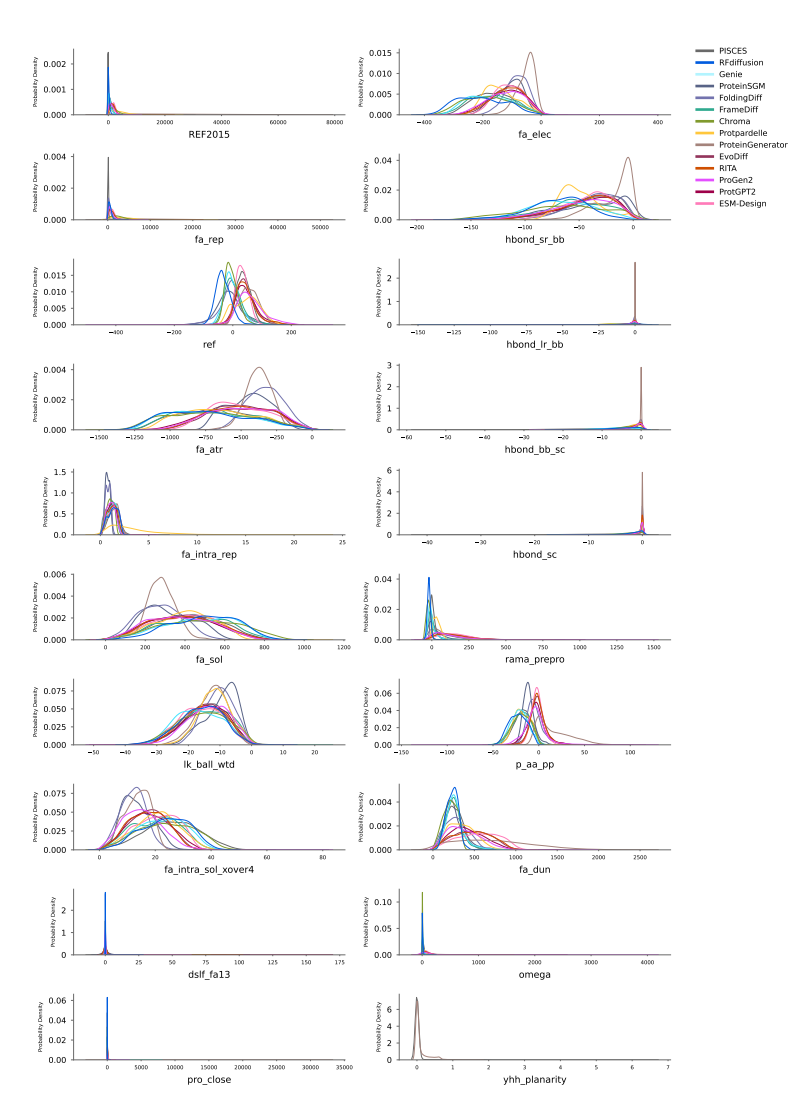
**Supplementary Figure 5. *Probability density estimation (KDE) of Rosetta Energy Score (REF2015) values and subterms for our generated monomers and PISCES chains.***

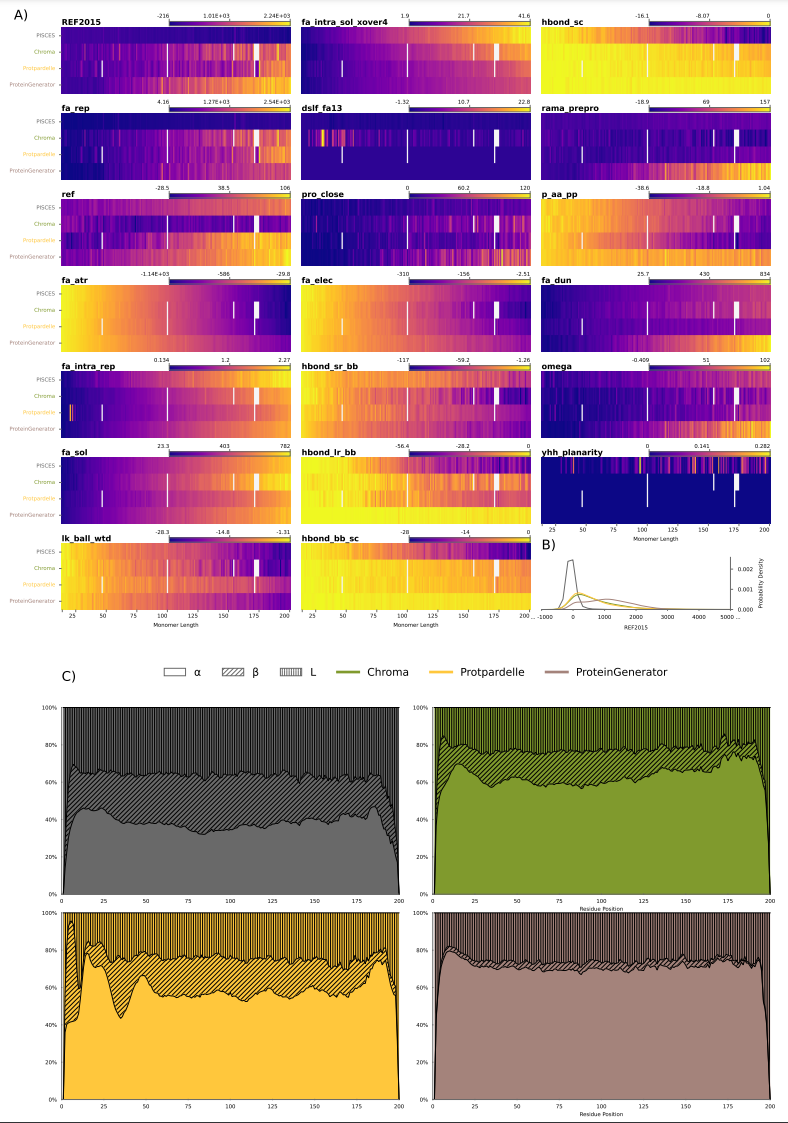

### **Supplementary Figure 6. *Rosetta Energy Scores (REF2015), subterm values, and secondary structure enrichment vs monomer length/position for PISCES chains and the structures predicted by OmegaFold for the primary sequences of monomer designs from Chroma, Protpardelle, and ProteinGenerator.***

*A) Energy Scores and subterm values vs monomer length. B) Probability density estimation (KDE) of REF2015 values. C) Percent (%) enrichment for alpha helix, beta strand and coil secondary structures per residue position.*

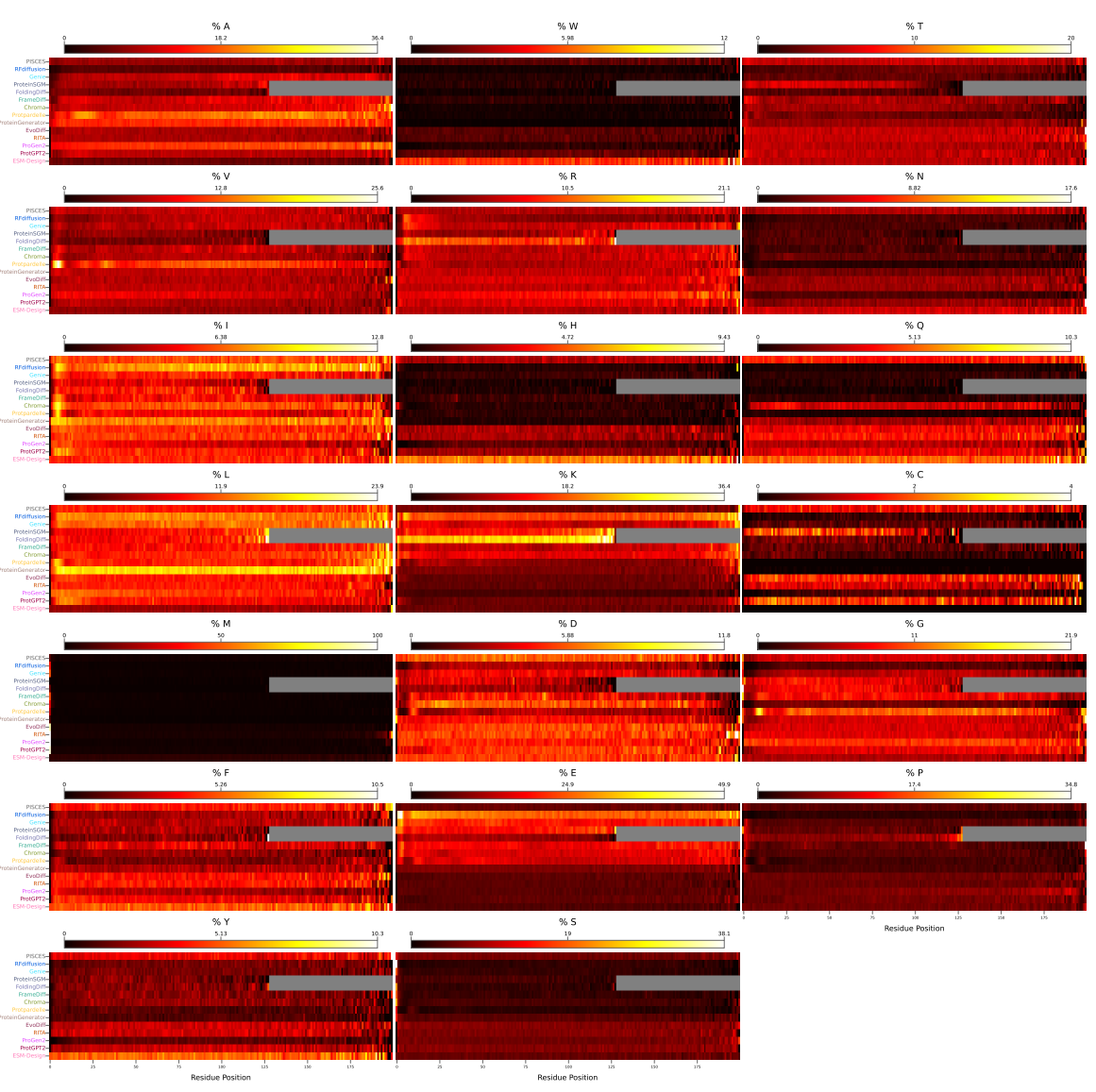

### **Supplementary Figure 7. *% Amino acid enrichment per position for our generated monomers and PISCES chains.***

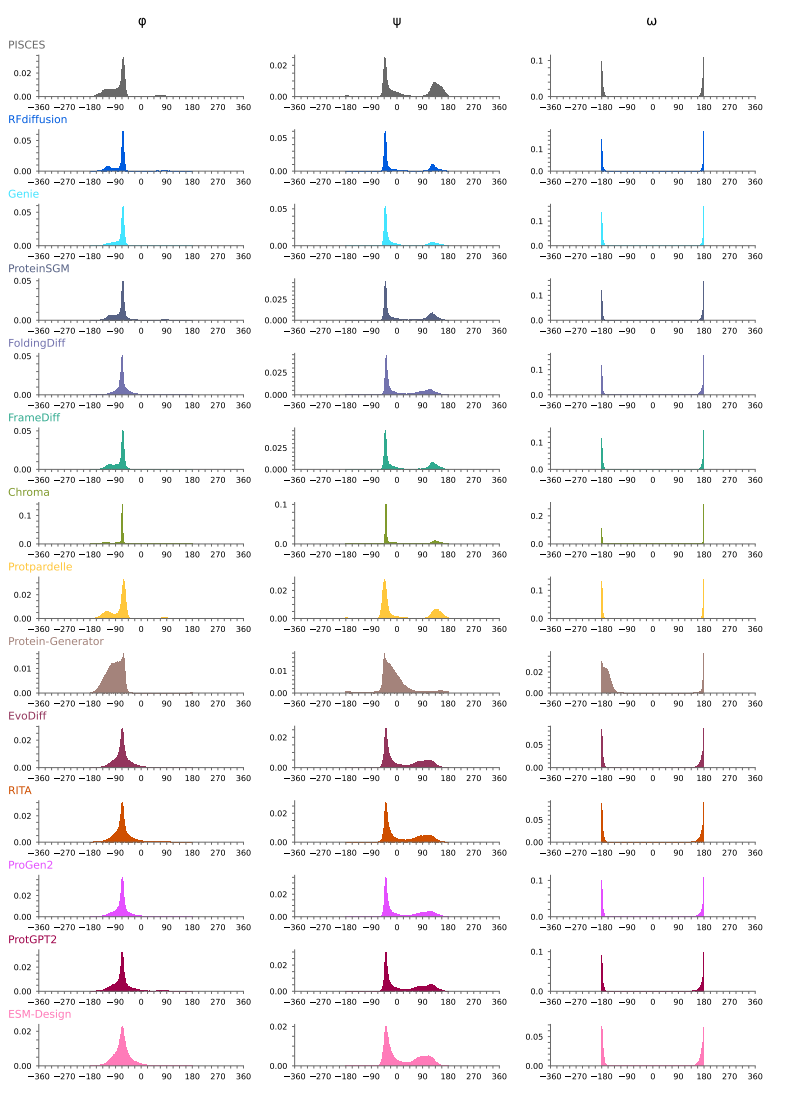

### **Supplementary Figure 8. *Distributions (KDE probability density estimation) of phi (Φ), psi (Ψ), and omega (ω) dihedral angles observed in our generated monomers and PISCES chains.***

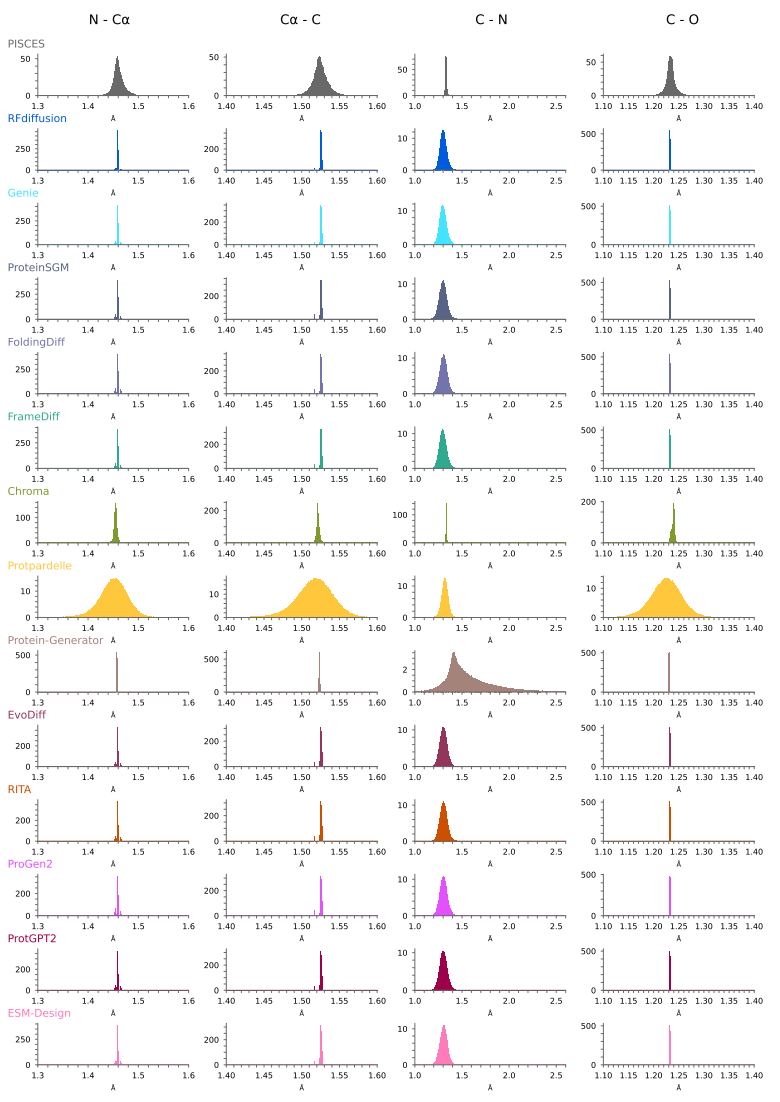

### **Supplementary Figure 9. *Distributions (KDE probability density estimation) of N-Cα, Cα-C, and C-O bond lengths observed in our generated monomers and PISCES chains.***

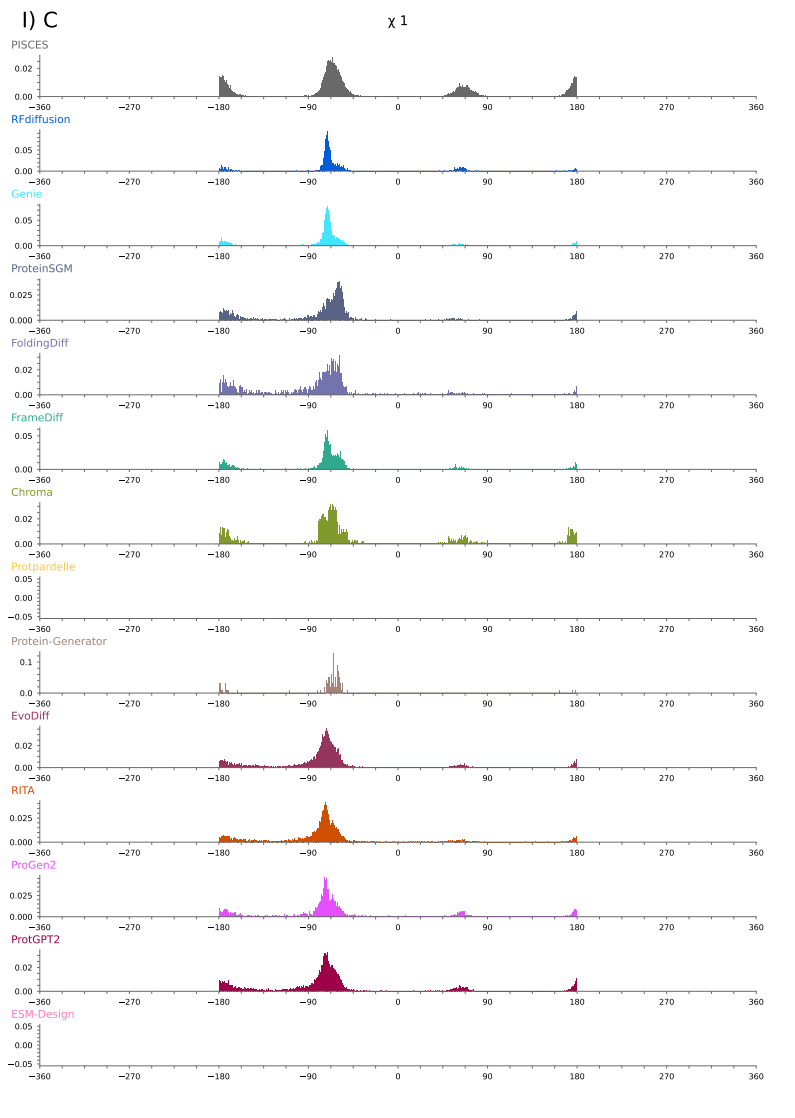

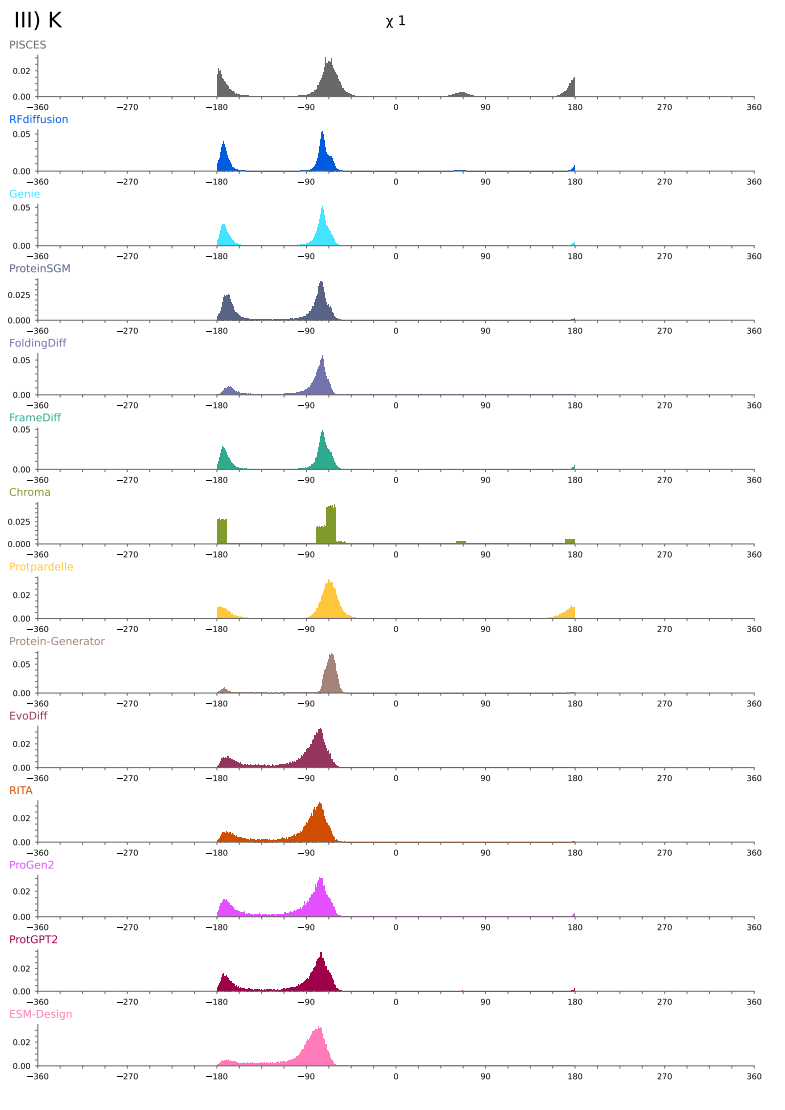

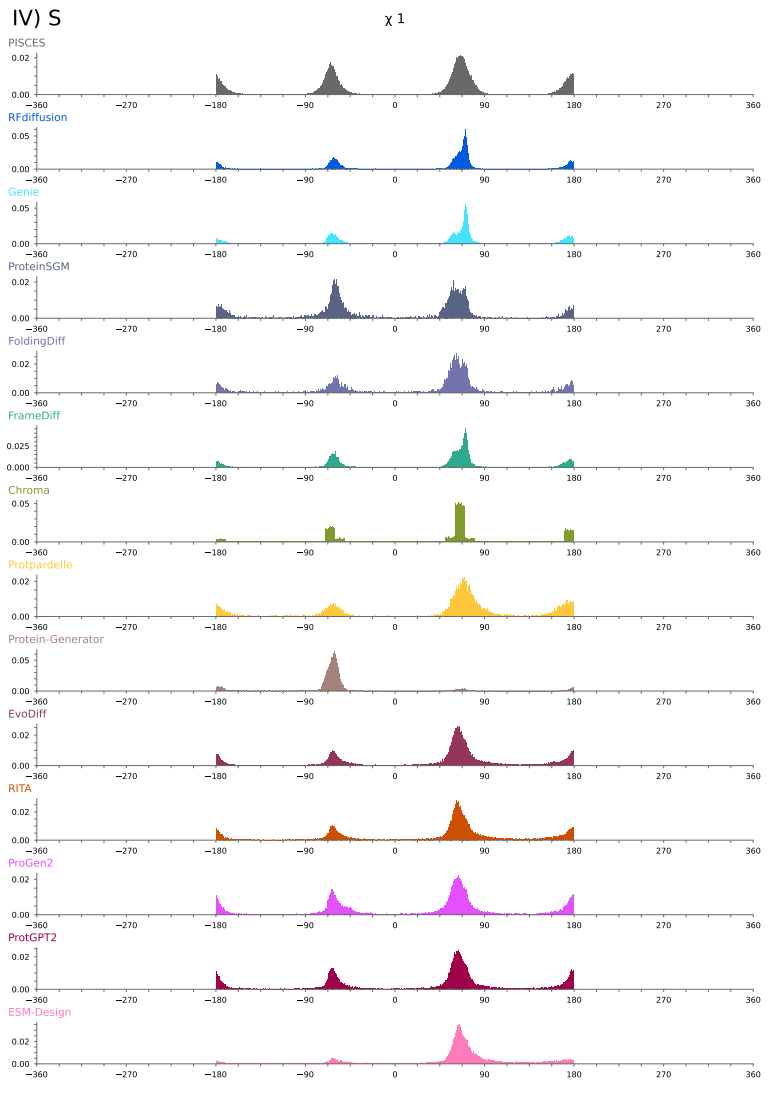

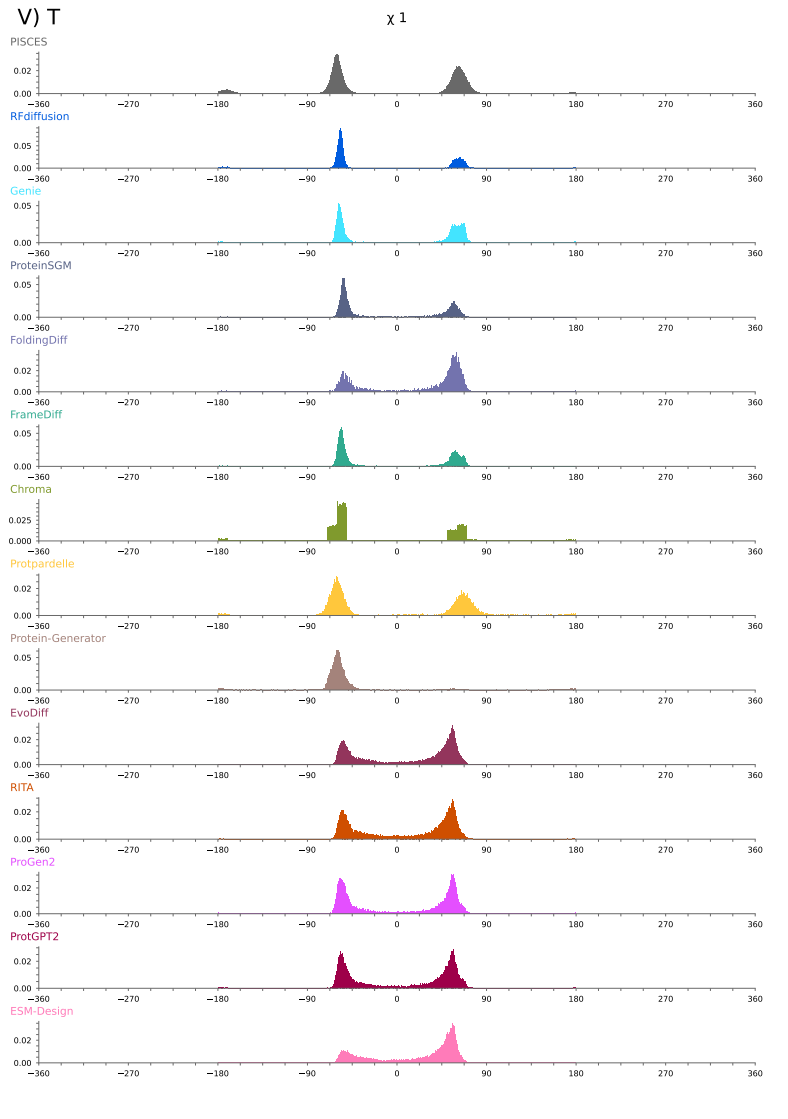

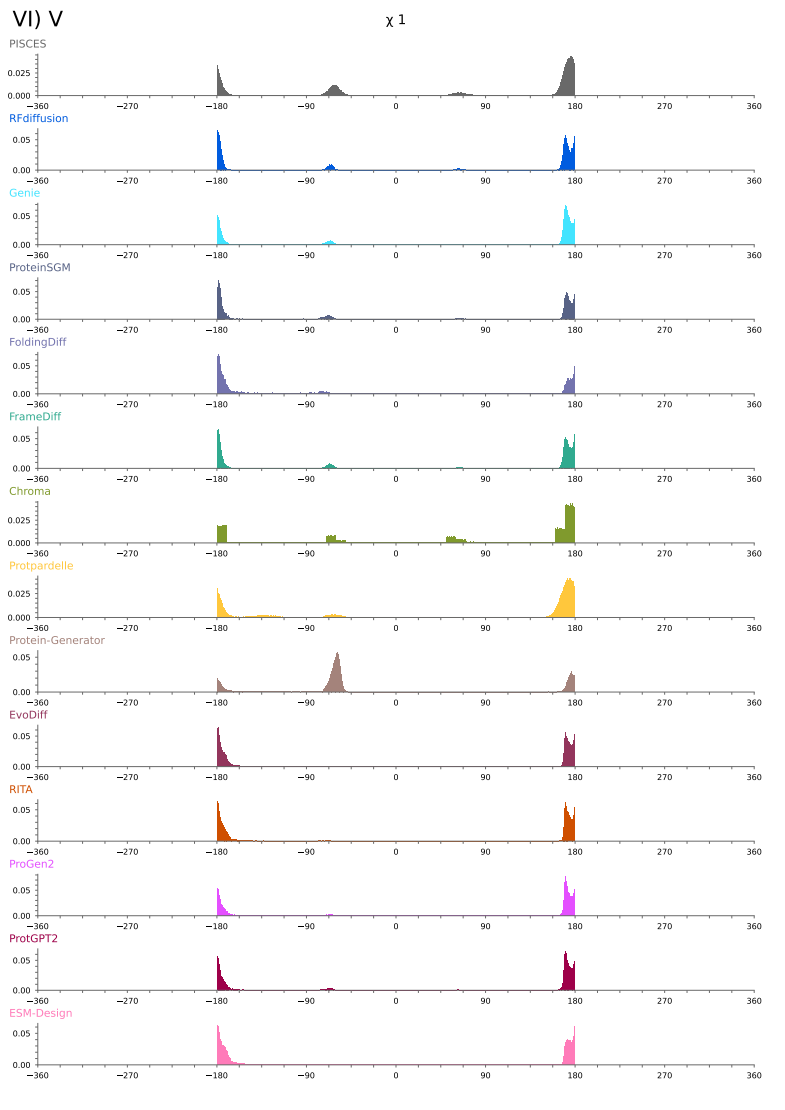

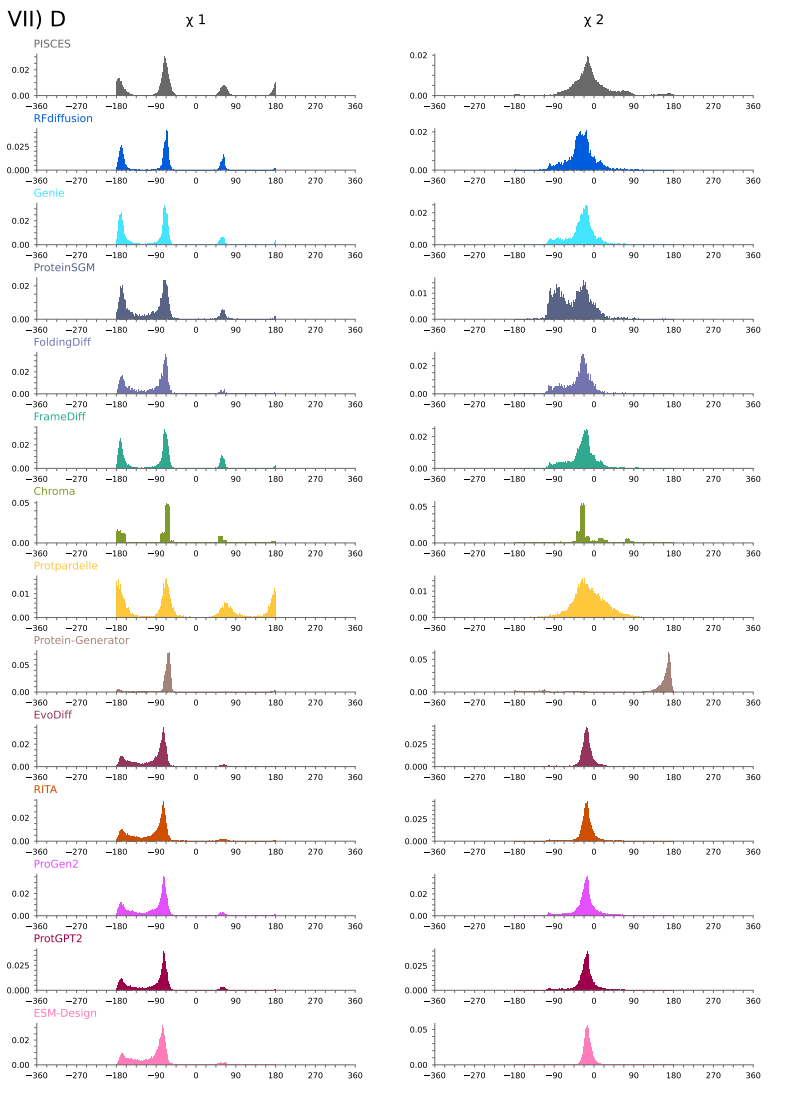

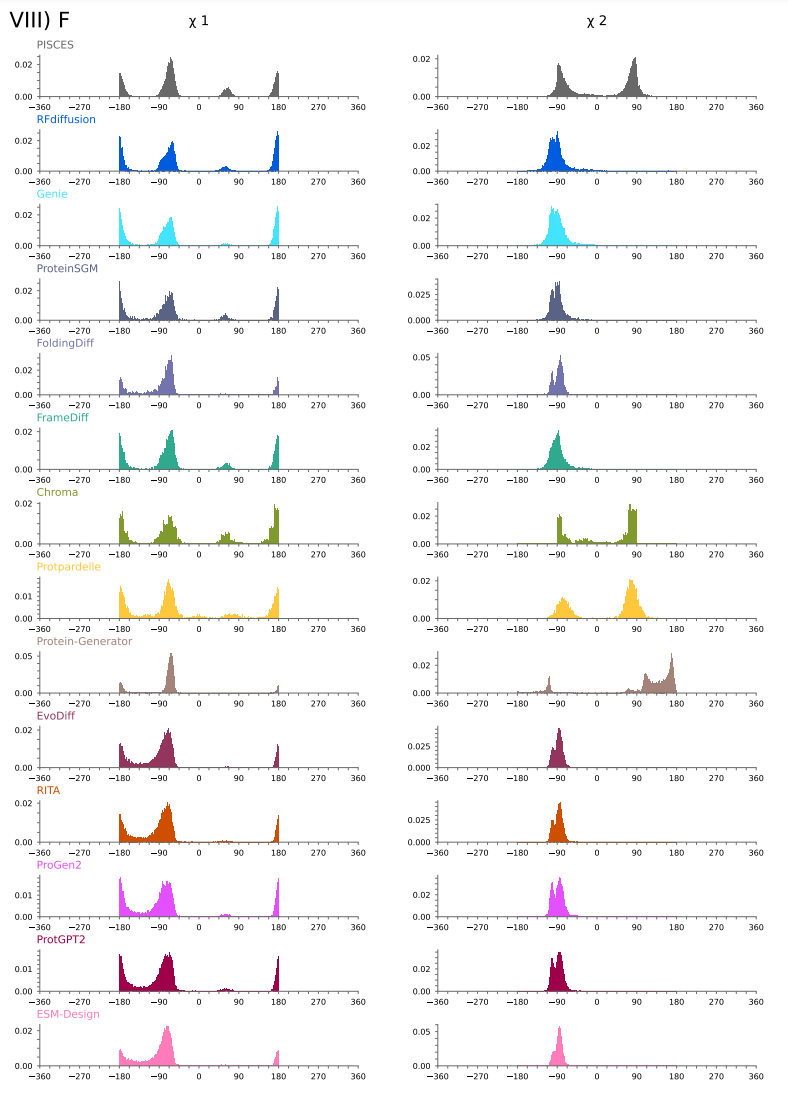

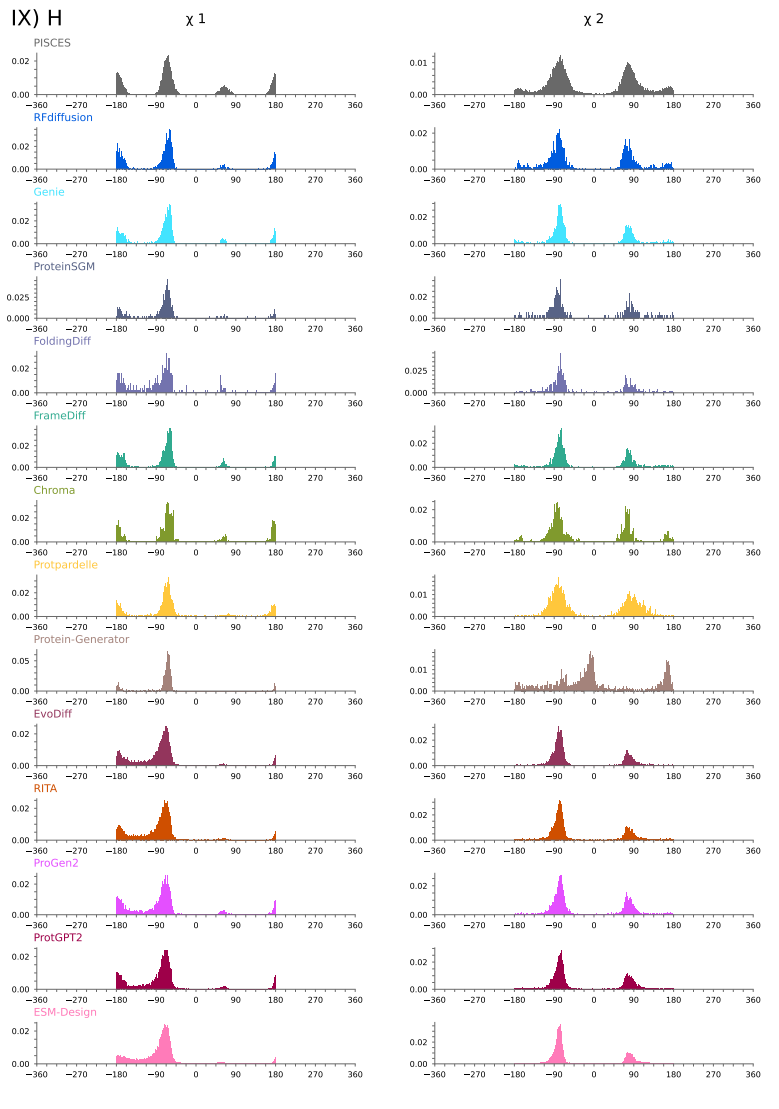

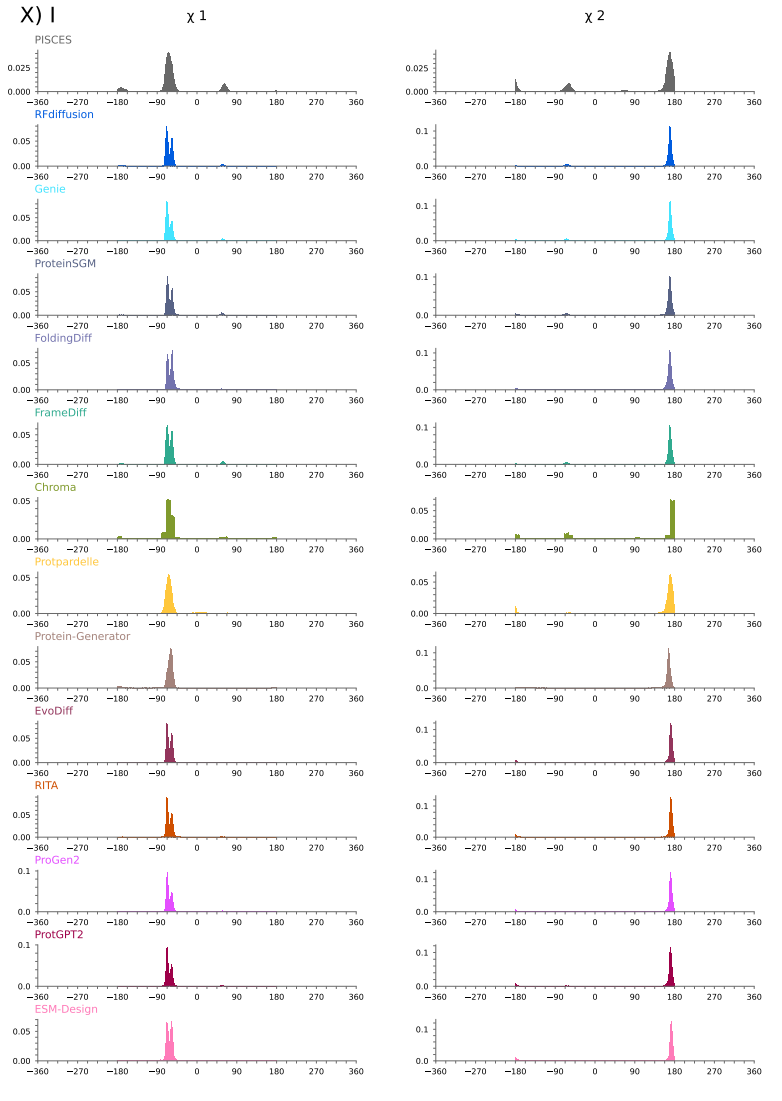

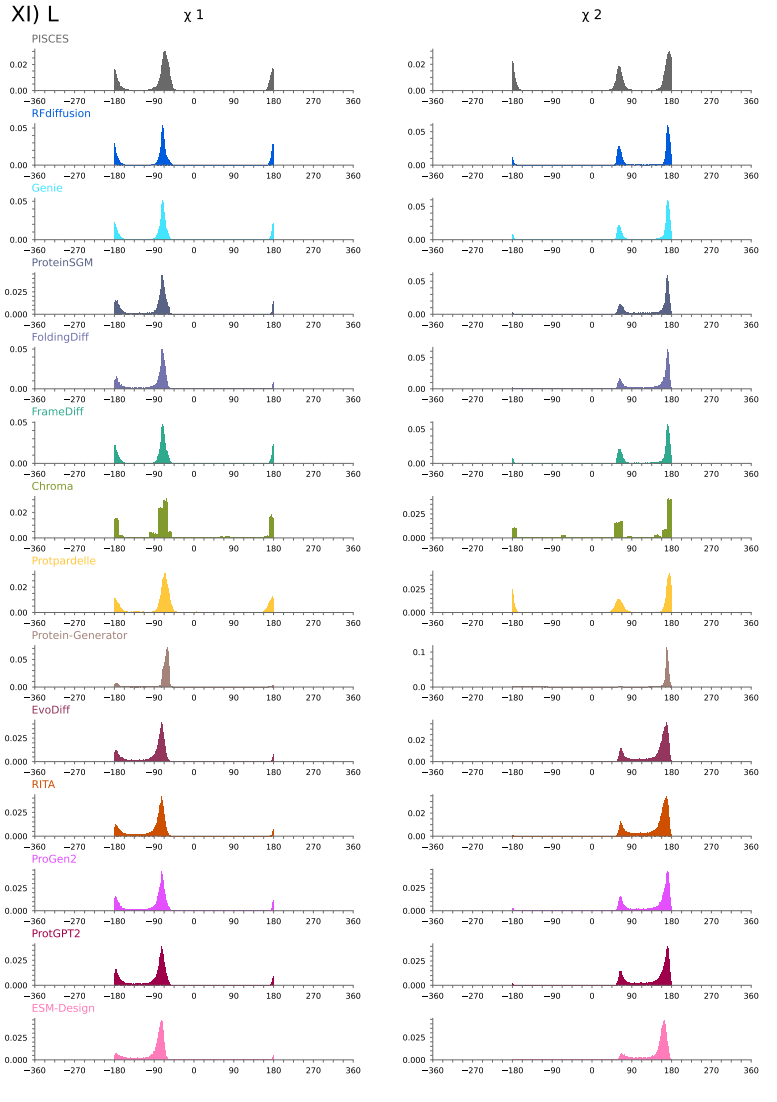

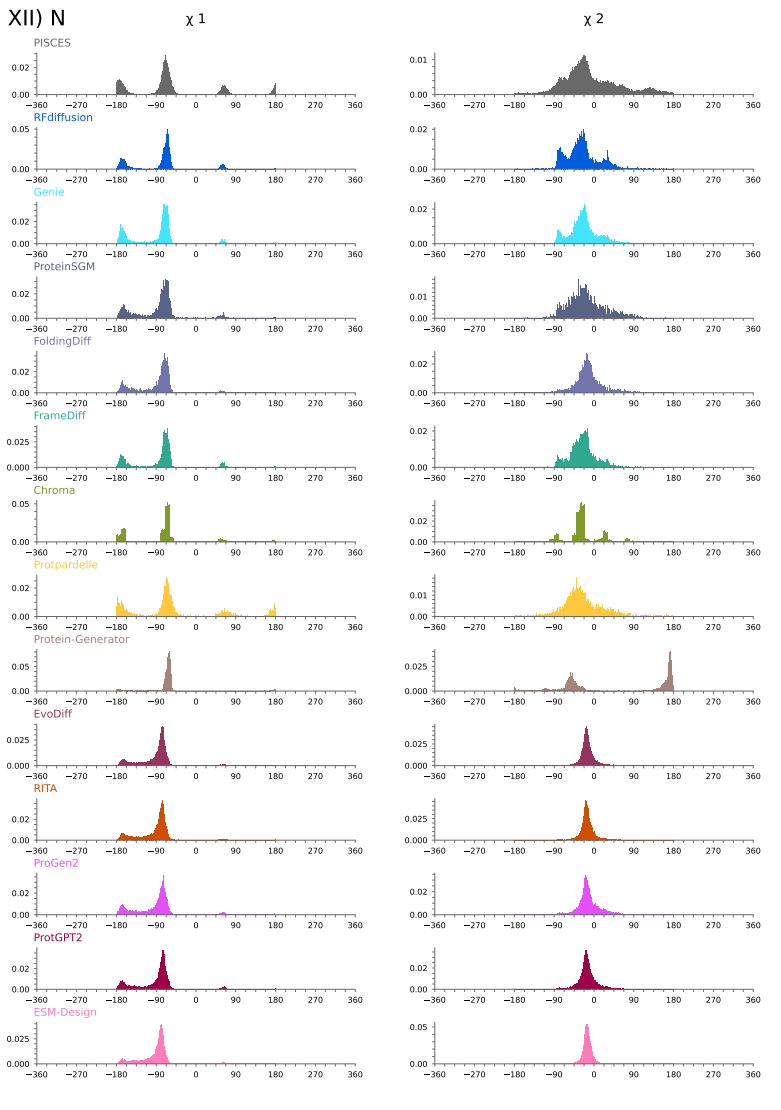

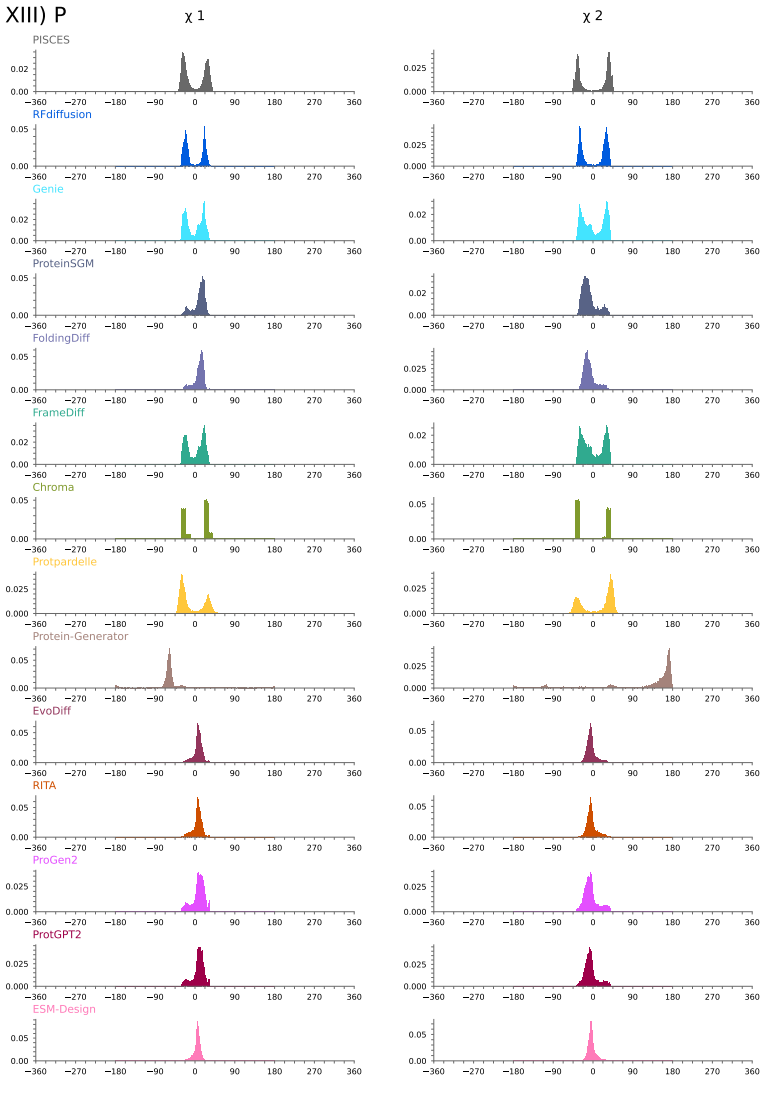

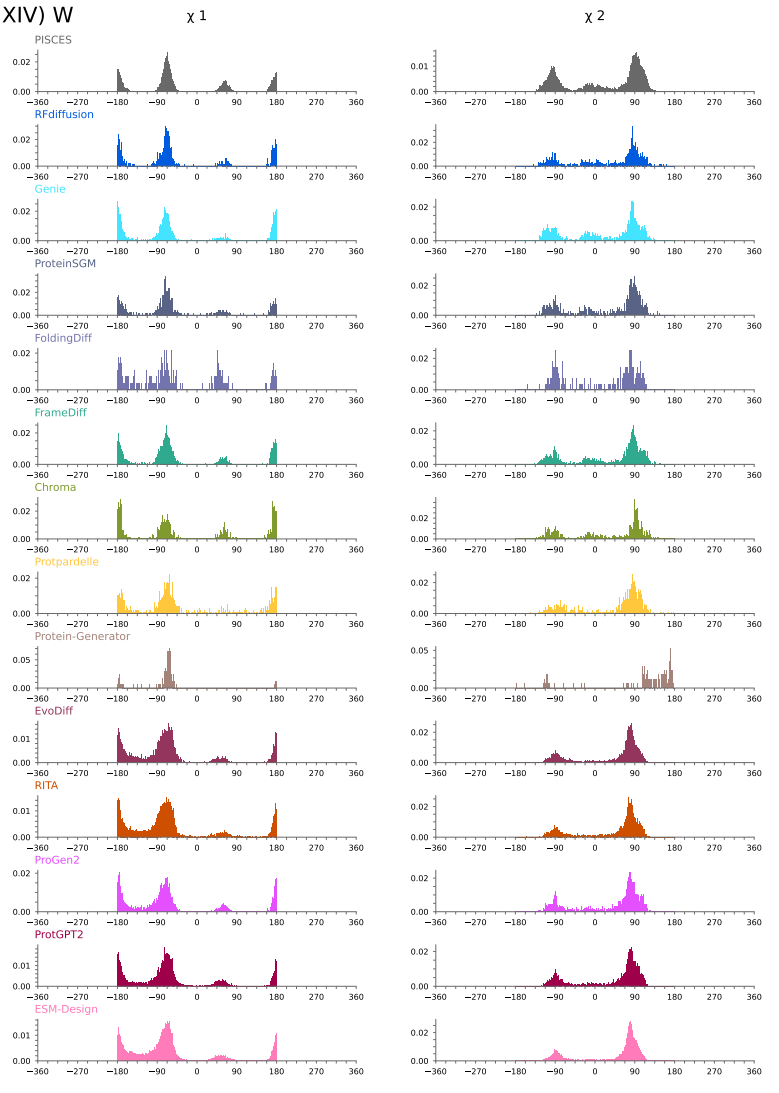

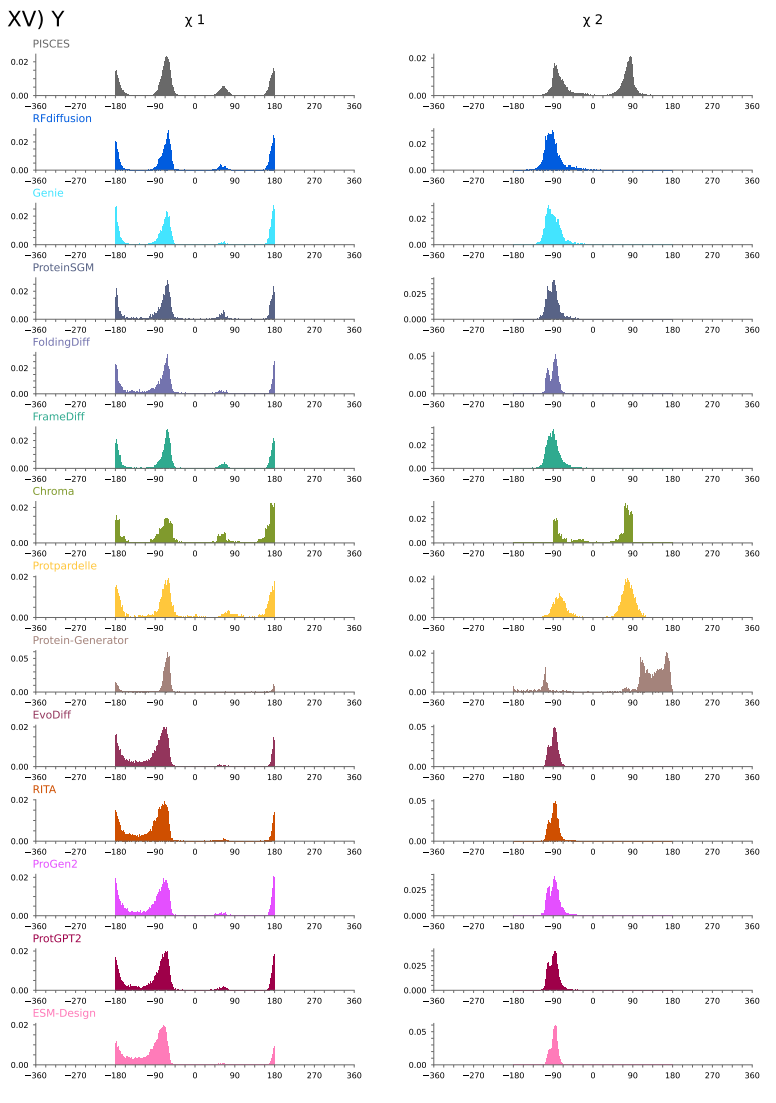

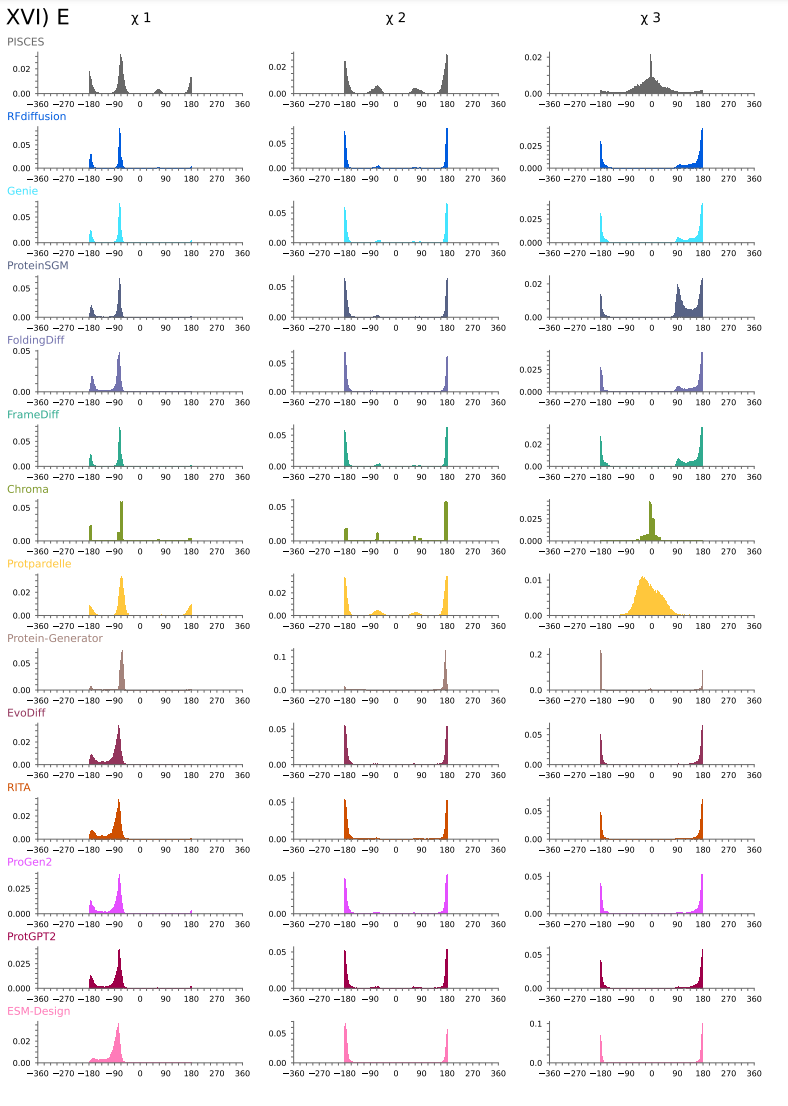

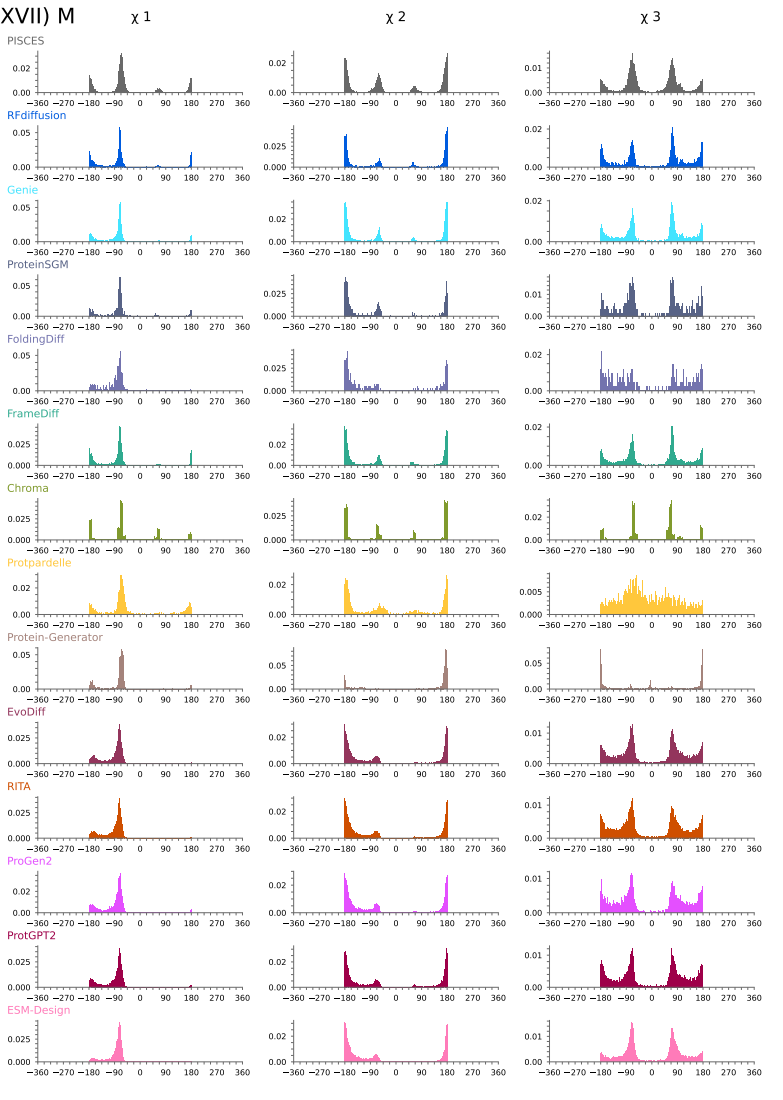

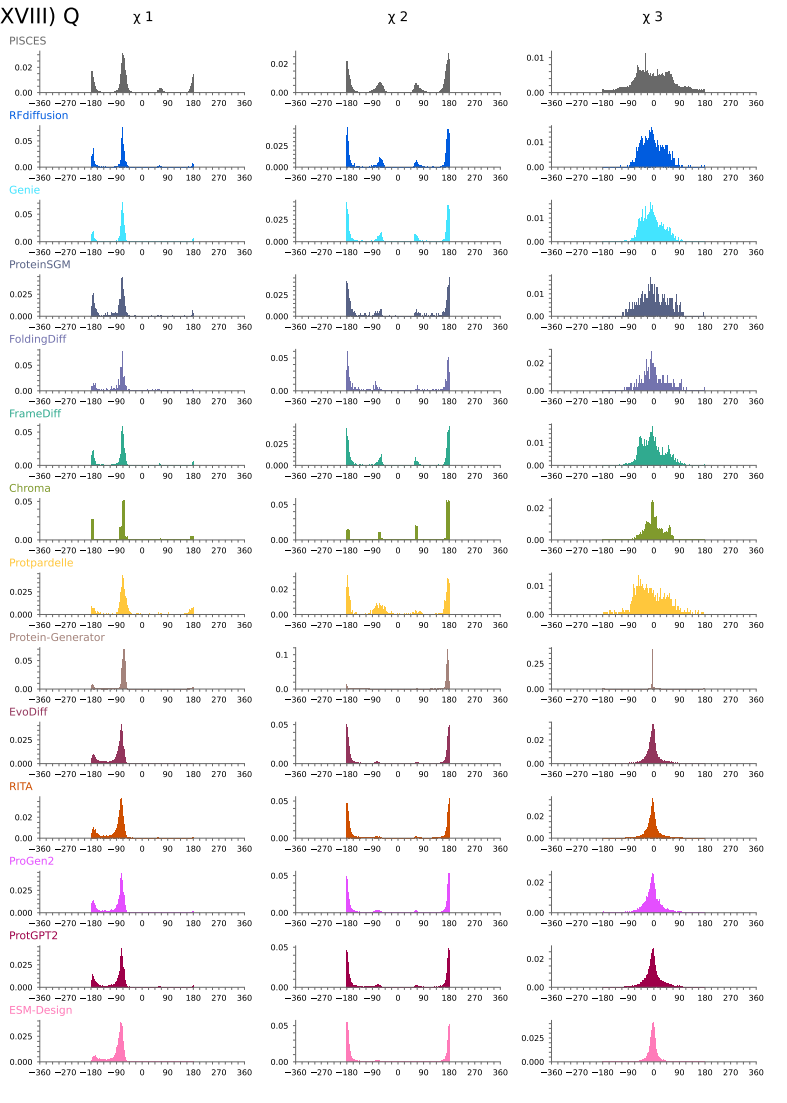

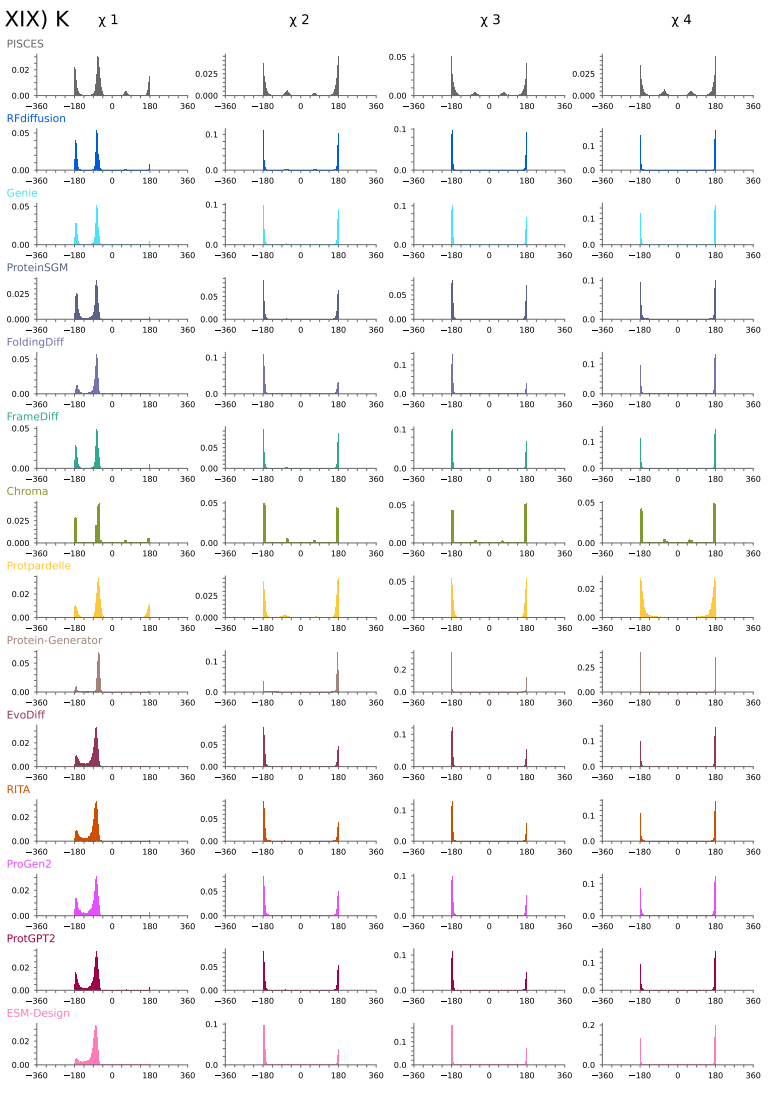

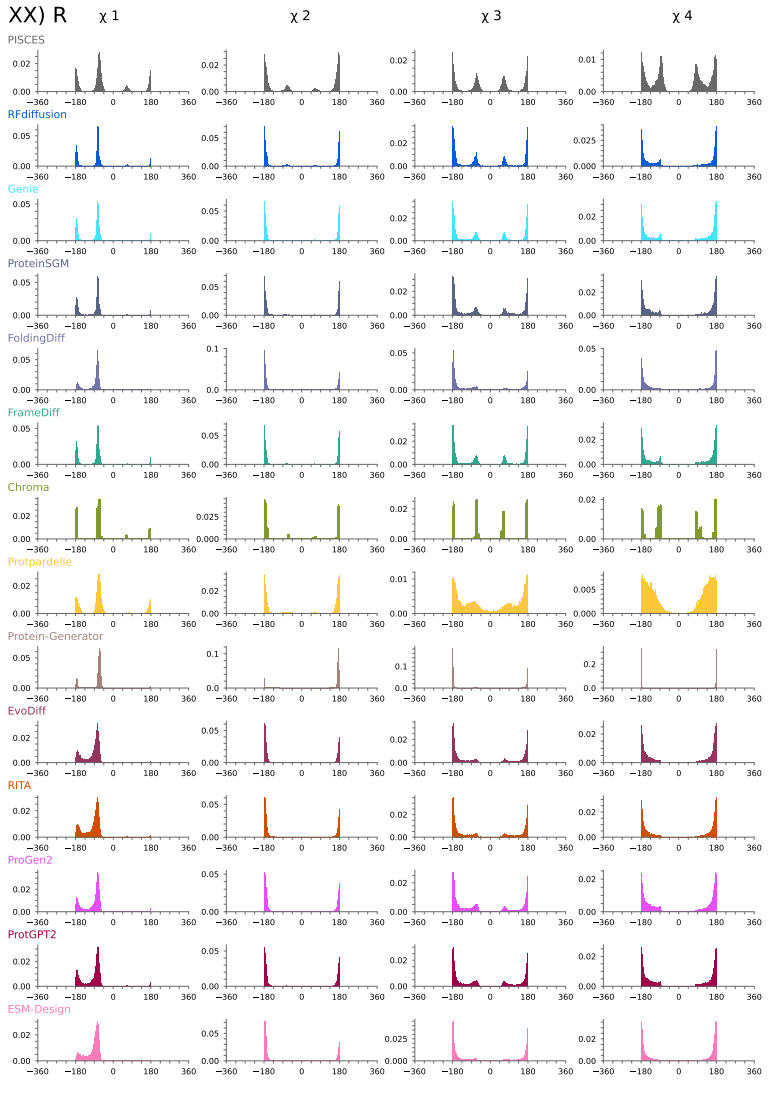

### **Supplementary Figure 10. *Distributions (KDE probability density estimation) of side-chain torsion angles ( χ_1_, χ_2_, χ_3_, χ_4_) observed in our generated monomers and PISCES chains for each residue type (i-XX).***

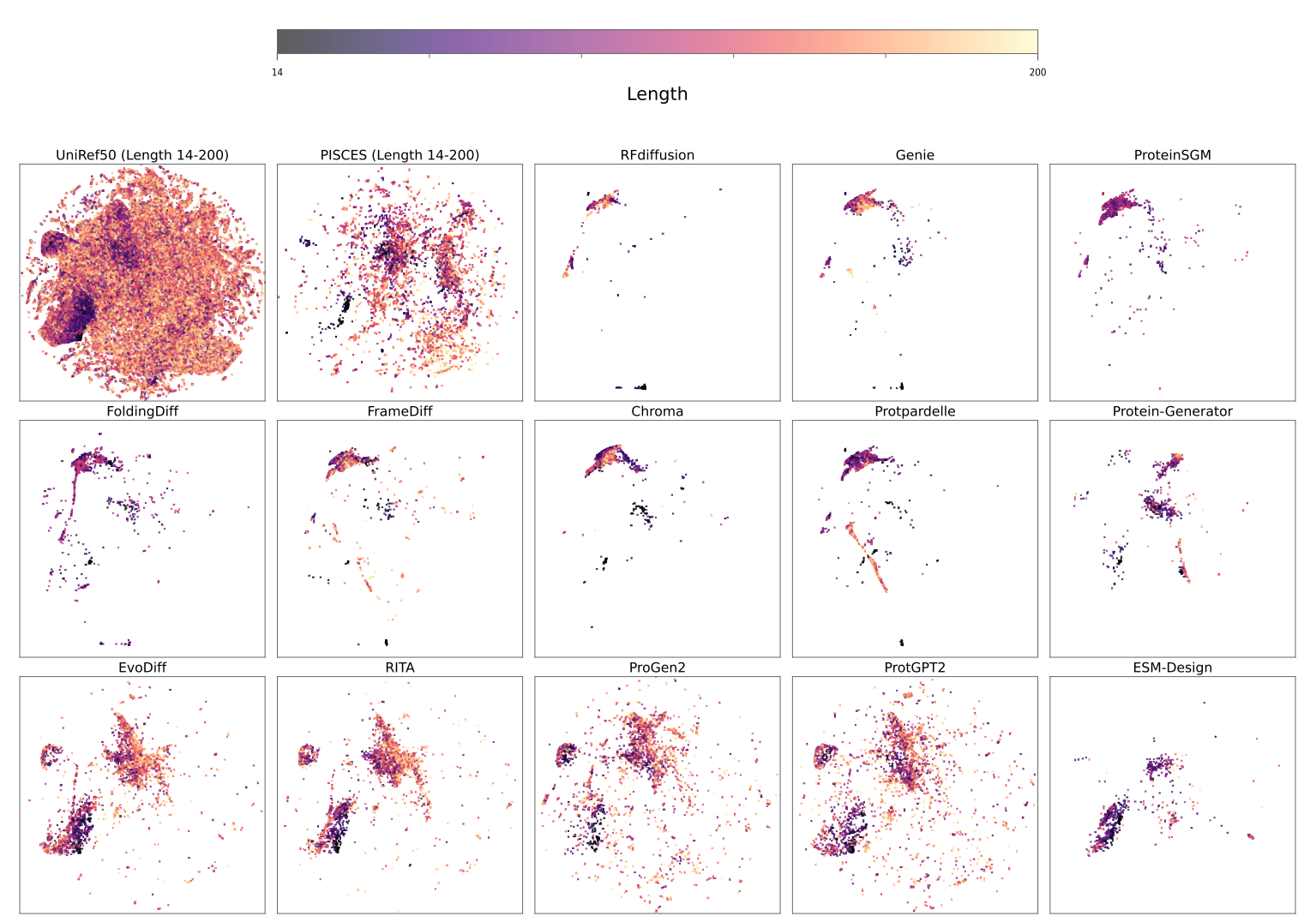

### **Supplementary Figure 11. *Distributions of sequence lengths of a 1% subsample of UniRef50, PISCES chains, and our generated monomers, throughout the t-SNE visualisations of their embedding-vectors in the ESM Large Language Model.***

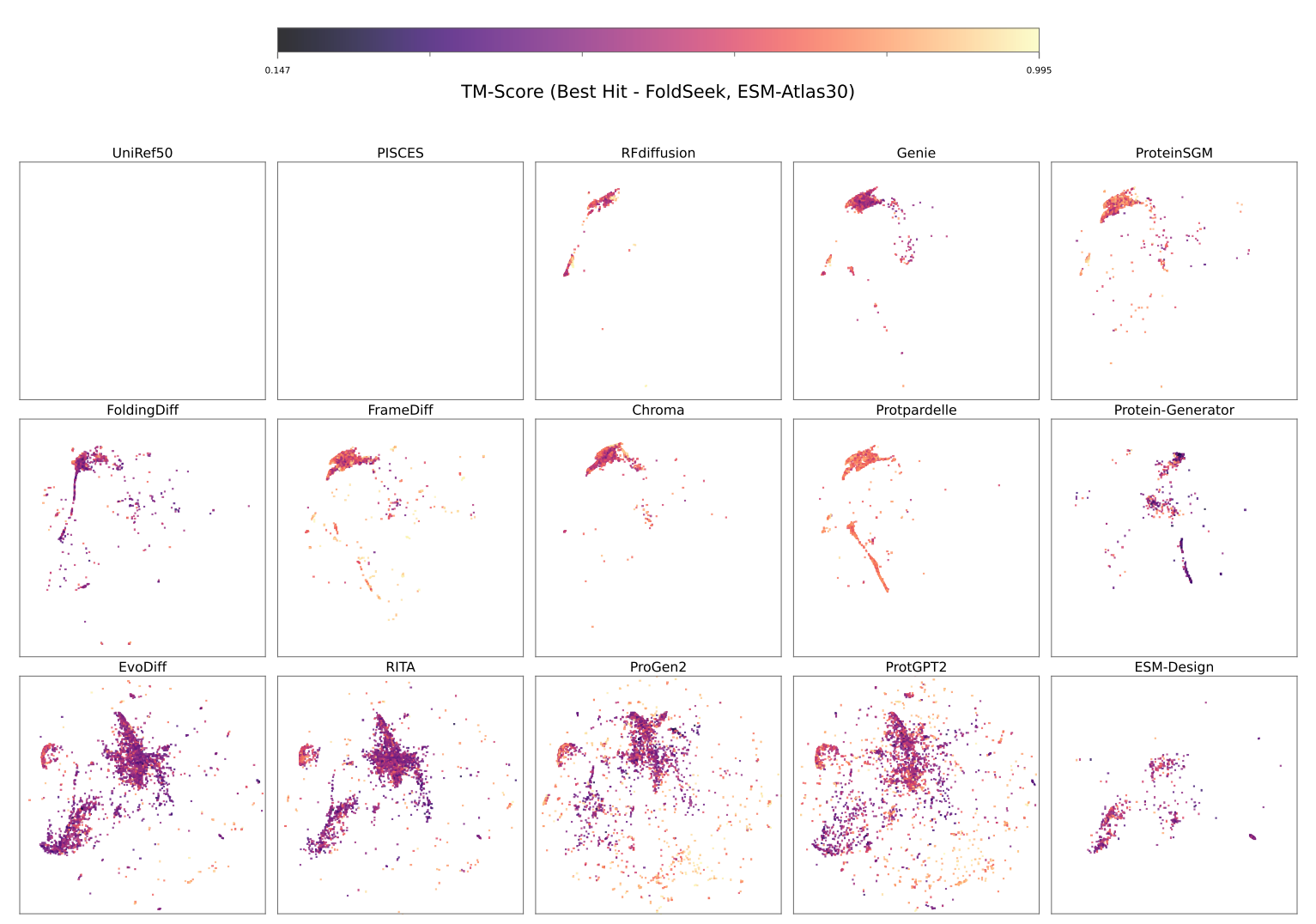

### **Supplementary Figure 12. *Distributions of the best-hit TM-Scores for our generated monomers when queried against the ESMAtlas30 database in the t-SNE visualisations of their embedding-vectors in the ESM Large Language Model.***

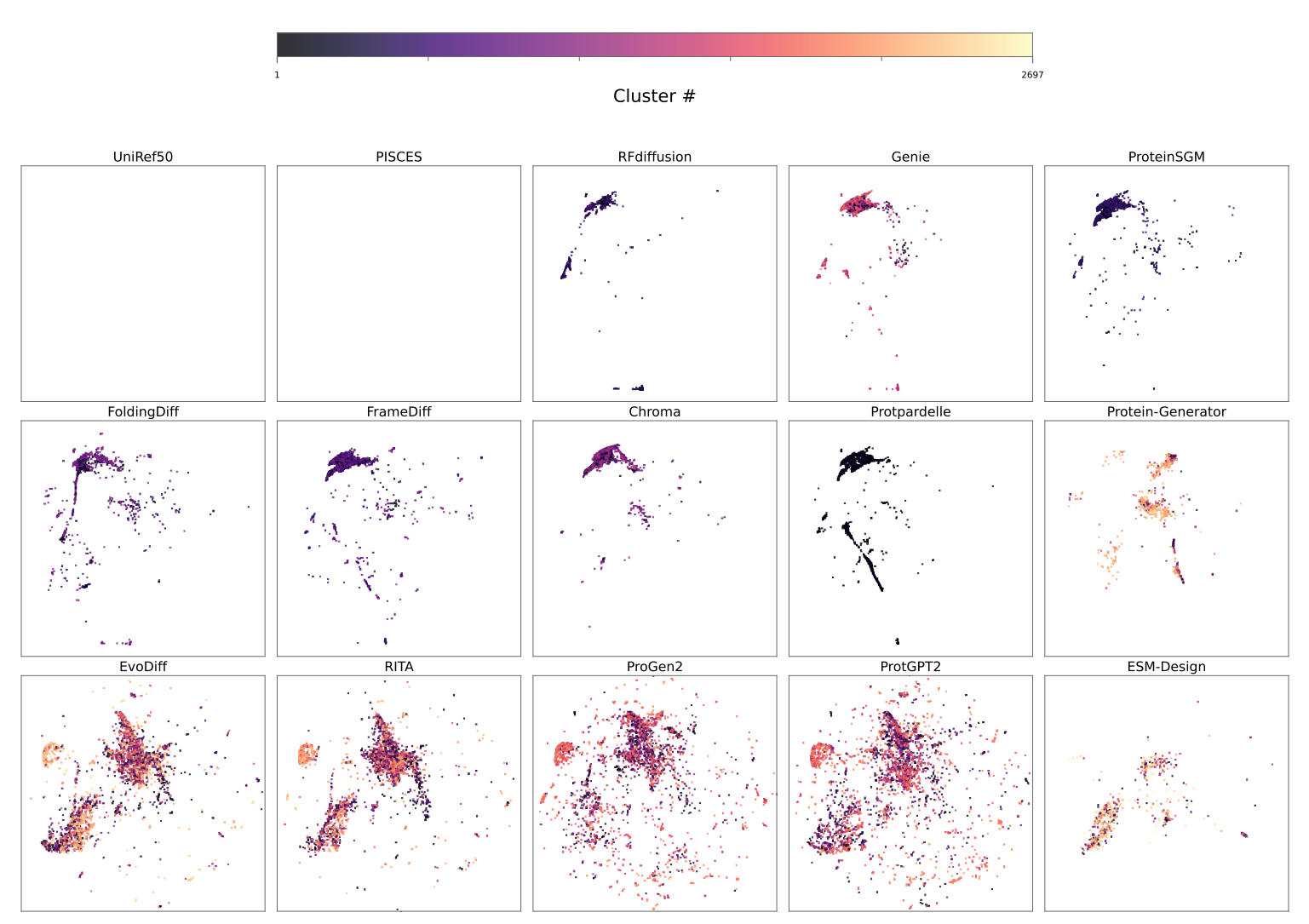

### **Supplementary Figure 13. *Distributions of the structural cluster labels identified by MaxCluster for our pools of generated monomers in the t-SNE visualisations of their embedding-vectors in the ESM Large Language Model.***

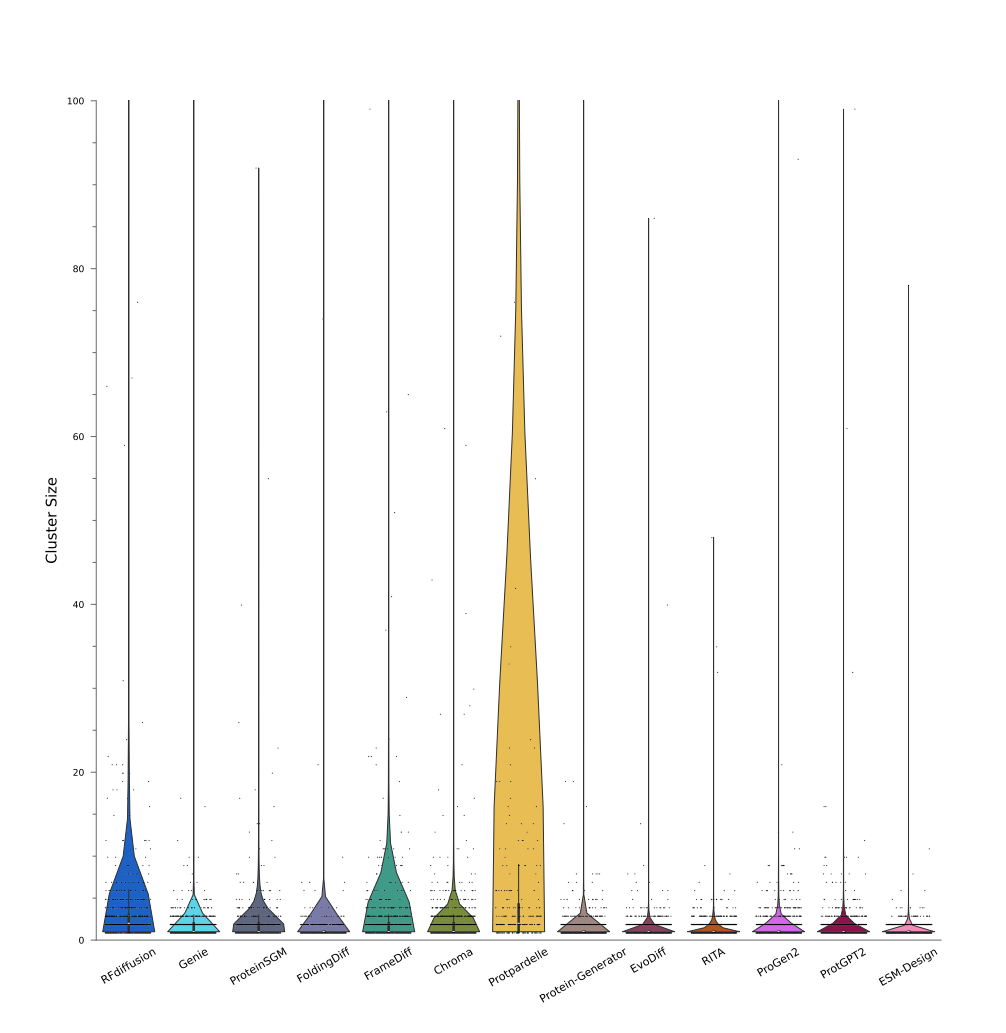

### **Supplementary Figure 14. *Distributions of sizes for structural clusters identified by MaxCluster for our pools of generated monomers.***

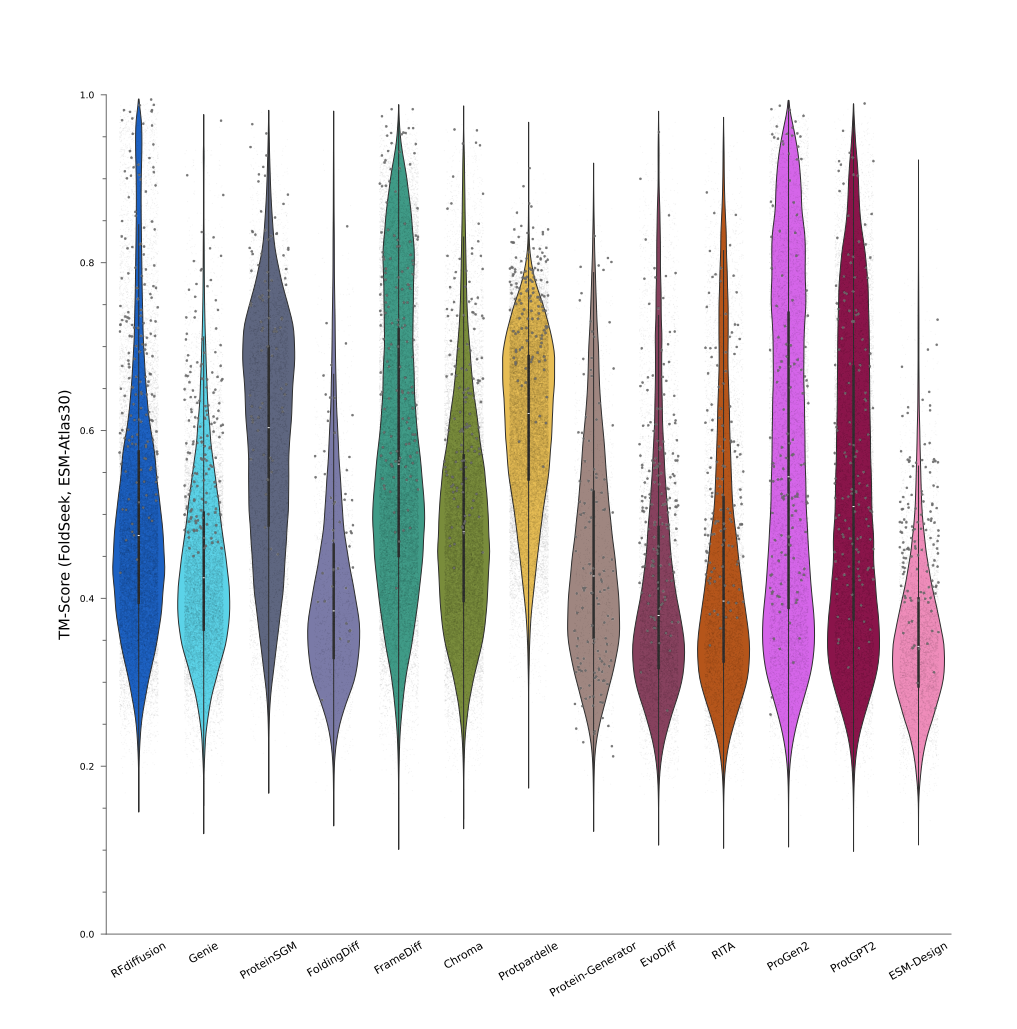

### **Supplementary Figure 15. *Distributions of TM-Scores of our generated monomers, by model, when queried against the ESMAtlas30 database.***

*Swarms represent ~5% of all hits. Larger swarm points represent TM-Scores from best hits.*
